# STIL overexpression shortens lifespan and reduces tumor formation in mice

**DOI:** 10.1101/2023.12.04.569842

**Authors:** Amira-Talaat Moussa, Marco R. Cosenza, Timothy Wohlfromm, Katharina Brobeil, Anthony Hill, Annarita Patrizi, Karin Müller-Decker, Tim Holland-Letz, Anna Jauch, Bianca Kraft, Alwin Krämer

## Abstract

Centrosomes are the major microtubule organizing centers of animal cells. Supernumerary centrosomes are a common feature of human tumors and associated with karyotype abnormalities and aggressive disease, but whether they are cause or consequence of cancer remains controversial. Here, we analyzed the consequences of centrosome amplification by generating transgenic mice in which centrosome numbers can be increased by overexpression of the structural centrosome protein STIL. We show that STIL overexpression induces centrosome amplification and aneuploidy, leading to senescence, apoptosis, and impaired proliferation in mouse embryonic fibroblasts, and microcephaly with increased perinatal lethality and shortened lifespan in mice. Importantly, both overall tumor formation in mice with constitutive, global STIL overexpression and chemical skin carcinogenesis in animals with inducible, skin-specific STIL overexpression were reduced, an effect that was not rescued by concomitant p53 inactivation. These results suggest that supernumerary centrosomes impair proliferation *in vitro* as well as *in vivo*, resulting in reduced lifespan and spontaneous as well as carcinogen-induced tumor formation.

## Introduction

Centrosomes are the major microtubule-organizing centers in mammalian cells and consist of a pair of centrioles embedded in pericentriolar material (Gonczy 2015). Centrosomes organize the bipolar spindle that partitions chromosomes during mitosis. Although centrosome amplification is associated with a growth disadvantage in cell lines (Holland et al. 2012; Sala et al. 2020), centrosome abnormalities are observed in the vast majority of human malignancies, where their presence correlates with karyotype abnormalities and poor prognosis. Aberrant centrosomes are found already in early stages of tumor development and lead to mitotic aberrations, which can cause chromosome missegregation in cultured cells (Ganem et al. 2009; Silkworth et al. 2009; Cosenza et al. 2017).

Centrosome duplication is controlled by the protein kinase PLK4 and the two structural centriole proteins STIL and SAS6 (Bettencourt-Dias et al. 2005; Habedanck et al. 2005; Strnad et al. 2007; Tang et al. 2011; Arquint et al. 2012; Vulprecht et al. 2012). Depletion of any one of these proteins blocks centrosome duplication and, conversely, overexpression causes centrosome amplification. Induction of extra centrosomes by overexpression of PLK4, the principal kinase regulating centrosome duplication, caused tumorigenesis in several animal models *in vivo*. In flies, overexpression of the PLK4 homolog SAK initiated tumors from larval brains with extra centrosomes in transplantation assays (Basto et al. 2008). In mice, centrosome amplification by inducible overexpression of PLK4 enhanced spontaneous tumor formation in multiple tissues in both p53-deficient and p53-wildtype backgrounds (Coelho et al. 2015; Levine et al. 2017; Shoshani et al. 2021). Although constitutive skin-specific PLK4 overexpression led to apoptosis of epidermal progenitors, skin barrier defects resulting in increased postnatal lethality, skin tumor formation was increased in the surviving mice with concomitant p53 deletion (Sercin et al. 2016). On the other hand, centrosome amplification by constitutive overexpression of PLK4 in embryonic neural progenitors resulted in microcephaly but did not promote tumorigenesis even after deletion of p53 (Marthiens et al. 2013). Similarly, neither global nor skin-specific induction of PLK4-driven centrosome amplification promoted spontaneous tumor formation or chemical skin carcinogenesis in mice, regardless of p53 status (Kulukian et al. 2015; Vitre et al. 2015).

Supernumerary centrosomes are believed to contribute to tumor formation via induction of chromosome missegregation and chromosomal instability (CIN). Whereas low CIN rates can be advantageous to tumor cells and support tumor formation and progression, high rates of CIN cause cell death and tumor suppression (Kops et al. 2004; Weaver et al. 2007; Funk et al. 2021). Accordingly, high-level centrosome amplification can cause cell death as a consequence of multipolar mitotic divisions *in vitro* and *in vivo*, especially in cells with inefficient centrosome clustering mechanisms (Ganem et al. 2009; Silkworth et al. 2009; Marthiens et al. 2013; Sercin et al. 2016). Differences in the extent of supernumerary centrosomes might therefore have contributed to the inconsistent results on tumor formation of the PLK4 mouse models. Also, PLK4 kinase has additional substrates and roles outside the centrosome and mitosis, which might impact on tumor development as well (Martindill et al. 2007; Rosario et al. 2010; Puklowski et al. 2011; Kazazian et al. 2017).

In addition to its impact on tumorigenesis, high levels of CIN, induced by hypomorphic alleles of the spindle assembly checkpoint protein BubR1, are associated with increased senescence and various progeroid and age-related phenotypes, including short lifespan, in mice (Baker et al. 2004). Remarkably, despite having severe aneuploidy, BubR1 hypomorphic animals do not have an increased spontaneous tumor burden. Similarly, Bub3/Rae1 haploinsufficient mice display a reduced lifespan but no increased spontaneous tumorigenesis either, despite accumulation of substantial aneuploidy (Baker et al. 2006).

To assess the impact of centrosome amplification on CIN, senescence, lifespan and tumor formation *in vivo* without interfering with extracentrosomal traits, we generated transgenic mouse models overexpressing the structural centrosome protein STIL, a 1,288 amino acid protein that is recruited to the proximal end of the mother centriole after phosphorylation by PLK4 to mark the procentriole emergence site (Gonczy 2015). At metaphase-anaphase transition the cytoplasmic bulk of STIL is degraded via the anaphase-promoting complex/cyclosome (APC/C)-proteasome pathway (Arquint et al. 2012). STIL mutations associated with resistance to proteasomal degradation and centrosome amplification are a cause of primary microcephaly (Kumar et al. 2009; Marthiens et al. 2013). Overexpression of STIL mRNA and protein has been found in many cancer types (Erez et al. 2004; Wang et al. 2022). However, little is known about its role in tumorigenesis.

## Results

### STIL overexpression induces centrosome amplification and aneuploidy in vitro

To investigate the consequences of centrosome amplification *in vivo*, we crossed transgenic C57BL/6 mice conditionally overexpressing STIL (B6-STIL) to mice expressing CRE recombinase under control of the CMV promoter (CMV-CRE) (Supplemental Fig. S1), which leads to ubiquitous transgene expression at levels similar to the CAG promoter used in most of the mouse models overexpressing PLK4 (Marthiens et al. 2013; Kulukian et al. 2015; Vitre et al. 2015; Sercin et al. 2016). CMV-CRE is ubiquitously expressed from early embryogenesis onwards. To characterize the effect of STIL overexpression *in vitro*, primary MEFs were derived from B6-STIL control, CMV-STIL hemizygous (CMV-STIL^+/-^) and CMV-STIL homozygous (CMV-STIL^+/+^) embryos. As expected, a graded expression of the transgene was found on transcript and protein level in early passage (p3) CMV-STIL MEFs (Fig. 1A,B; Supplemental Fig. S2). STIL level elevation induced substantial centrosome amplification and abnormal mitoses in CMV-STIL MEFs. Rosette-like arrangements of procentrioles were frequently observed. Despite the graded increase in PLK4 expression, CMV-STIL^+/-^ and CMV-STIL^+/+^ MEFs exhibited a similar increase in supernumerary centrioles (Fig. 1C,D), in line with earlier findings suggesting that only a limited amount of excess PLK4 can access the centrosome at a given time (Holland et al. 2012). Similar to what has been described for MEFs overexpressing PLK4, CMV-STIL MEFs clustered their extra centrosomes into pseudo-bipolar spindles with high efficiency (Fig. 1E). Accordingly, the rate of respective mitotic aberrations was increased affecting about 80% of observed CMV-STIL cell divisions (Fig. 1F).

**Figure 1.**
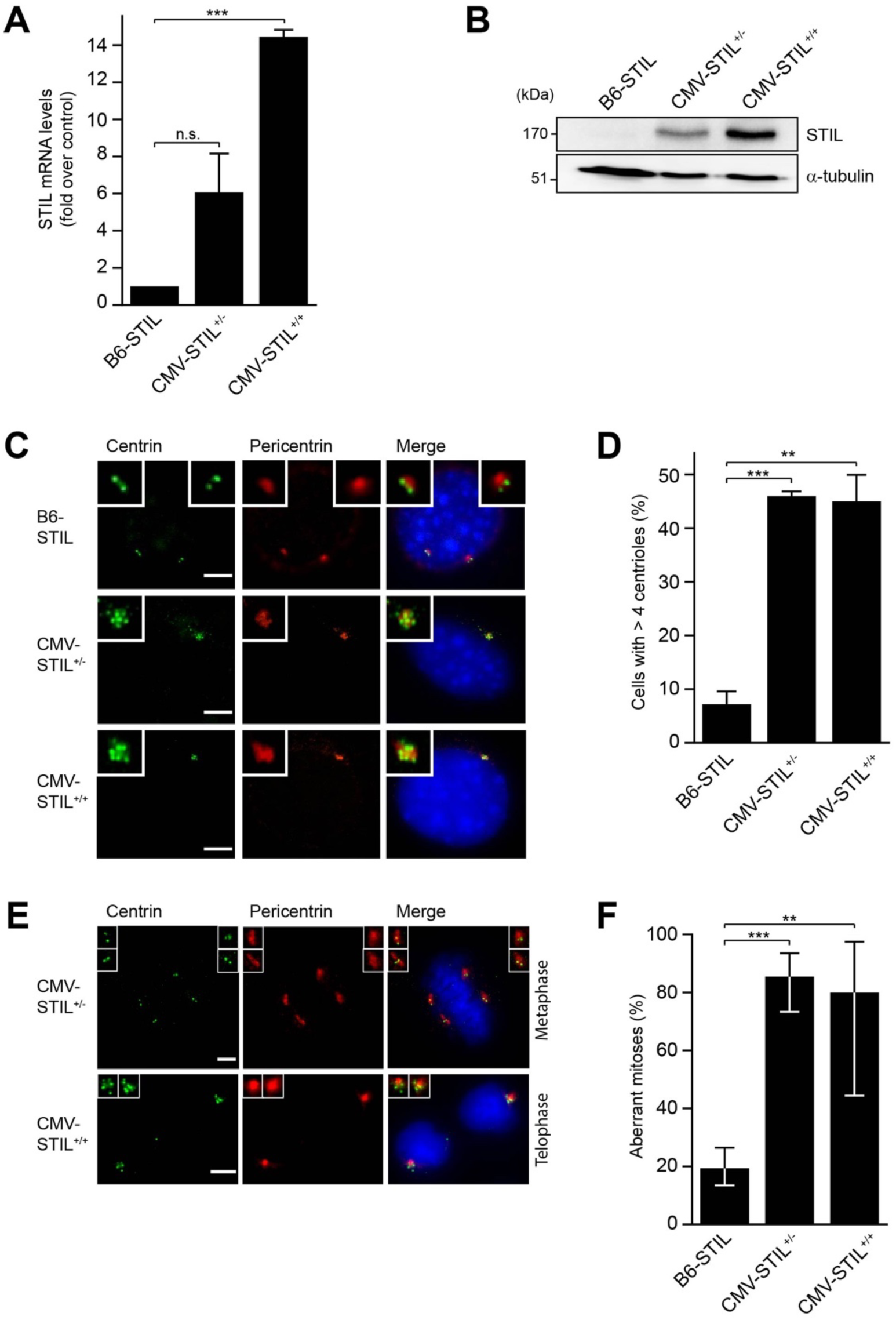
STIL overexpression induces centrosome amplification and aberrant mitoses *in vitro*. (*A*) Quantitative RT-PCR showing the fold increase of STIL mRNA levels in CMV-STIL^+/-^ and CMV-STIL^+/+^ compared to B6-STIL MEFs. Data are means ± SEM from three independent experiments. (*B*) Immunoblotting showing STIL protein expression levels in MEFs derived from B6-STIL, CMV-STIL^+/-^ and CMV-STIL^+/+^ mice, respectively. α-tubulin served as a loading control. (*C*) Immunofluorescence images of centrosomes in B6-STIL control (2 centrioles per centrosome), CMV-STIL^+/-^ and CMV-STIL^+/+^ MEFs (7 centrioles per centrosome each), immunostained with antibodies to centrin and pericentrin. DNA is stained with DAPI (blue). Centrosomes are shown enlarged in insets. Scale bar, 5 µm. (*D*) Percentage of B6-STIL control, CMV-STIL^+/-^ and CMV-STIL^+/+^ MEFs with supernumerary centrioles. Data are means ± SEM from three independent experiments. (*E*) Representative immunofluorescence images of metaphase CMV-STIL^+/-^ (upper panel; 2 centrosomes with 2 centrioles each per spindle pole) and telophase CMV-STIL^+/+^ (lower panel; 1 centrosome with 5 and 5 centrioles, respectively, per spindle pole) MEFs, immunostained with antibodies to centrin and pericentrin. DNA is stained with DAPI (blue). Centrosomes are shown enlarged in insets. Scale bar, 5 µm. (*F*) Percentage of mitotic B6-STIL control, CMV-STIL^+/-^ and CMV-STIL^+/+^ MEFs with aberrant mitoses. Data are means ± Clopper-Pearson 95%-CI. \**p*<0.05, \*\**p*<0.01, \*\*\**p*<0.001; n.s., not significant; Two-tailed Student’s *t*-tests were applied in (*A*) and (*D*), Fisher’s exact test in (*F*).

To determine whether these mitotic abnormalities lead to chromosome segregation errors and consequent aneuploidy, we assessed the frequencies of micronuclei and aberrant karyotypes in interphase and mitotic CMV-STIL MEFs, respectively. The frequency of micronuclei increased with STIL expression levels in CMV-STIL^+/-^ and CMV-STIL^+/+^ MEFs (*p*<0.001) (Fig. 2A,B). Similarly, multiplex fluorescence *in situ* hybridization (M-FISH) revealed a graded increase of metaphases with numerical and structural karyotype aberrations (*p*<0.01) and tetraploidization (*p*<0.01) (Fig. 2C-F), with more than half of the abnormal metaphases containing multiple karyotype aberrations in both CMV-STIL^+/-^ and CMV-STIL^+/+^ MEFs.

**Figure 2.**
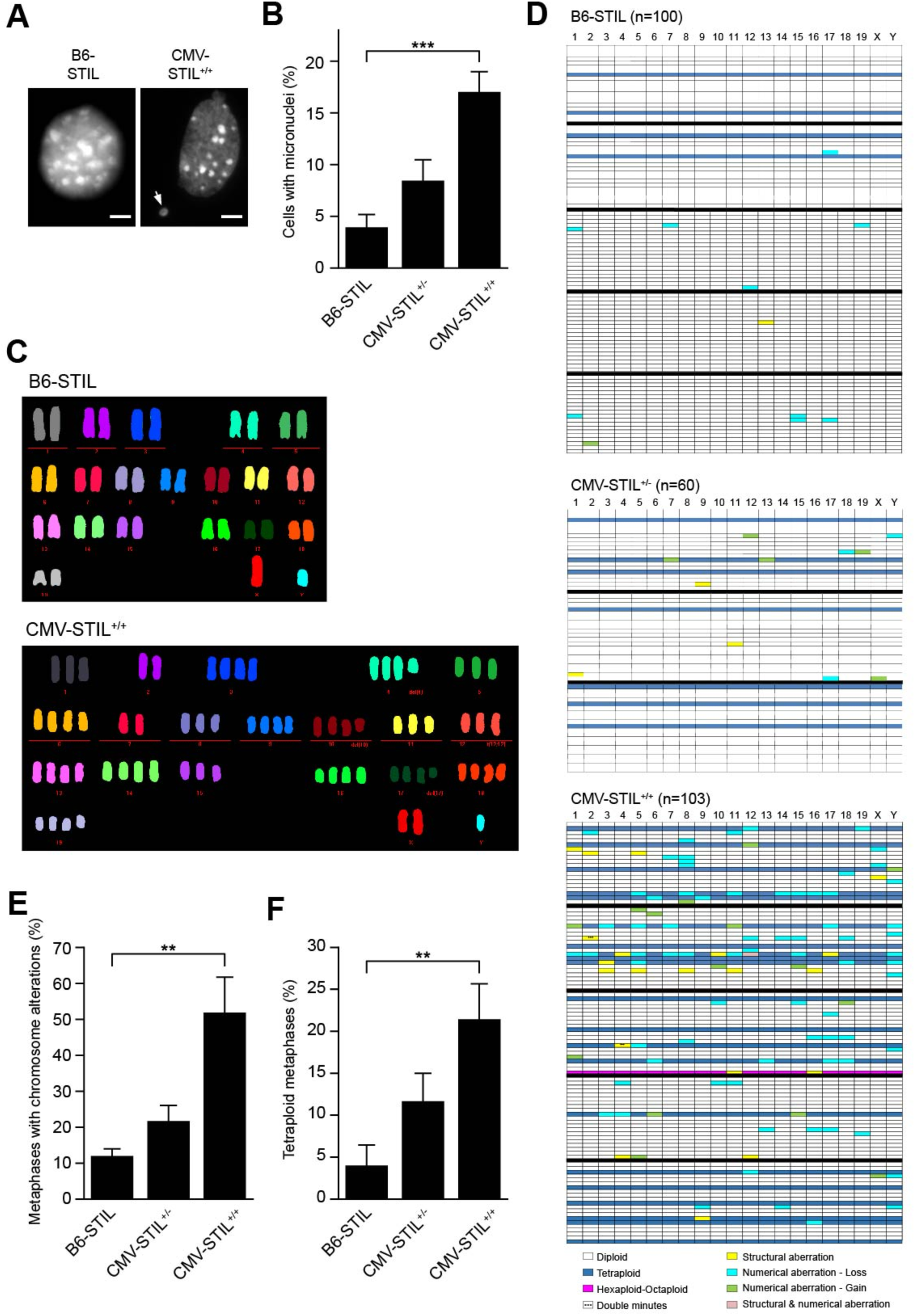
STIL overexpression induces aneuploidy *in vitro*. (*A*) Immunofluorescence images of nuclei in B6-STIL control and CMV-STIL^+/+^ MEFs. DNA is stained with Hoechst 33342. A micronucleus is marked with an arrow. Scale bar, 5 µm. (*B*) Percentage of B6-STIL control, CMV-STIL^+/-^ and CMV-STIL^+/+^ MEFs with micronuclei. Data are means ± SEM from three independent experiments. (*C*) Representative M-FISH metaphases of B6-STIL control and CMV-STIL^+/+^ MEFs. (*D*) Chromosome aberration profiles of B6-STIL control (n=100), CMV-STIL^+/-^ (n=60) and CMV-STIL^+/+^ (n=103) MEFs. Each row represents a single multicolor karyotyped metaphase, with chromosomes plotted as columns. Different colors are used to depict chromosome aberration types. Percentage of B6-STIL control, CMV-STIL^+/-^ and CMV-STIL^+/+^ MEF metaphases with chromosome aberrations (*E*) and tetraploid chromosome content (*F*) as calculated from (*D*). Data are means ± SEM. \**p*<0.05, \*\**p*<0.01, \*\*\**p*<0.001; two-tailed Student’s t-test.

### STIL overexpression impairs proliferation, and induces apoptosis and senescence in vitro

Centrosome amplification is associated with a growth disadvantage in cell lines and PLK4-overexpressing MEFs (Holland et al. 2012; Levine et al. 2017; Sala et al. 2020). Consistently, extra centrosomes led to a graded inhibition of the proliferation in CMV-STIL^+/-^ and CMV-STIL^+/+^ MEFs (Fig. 3A). Different from PLK4-overexpressing MEFs, where interference with p53 function alleviates the proliferation arrest (Levine et al. 2017), crossbreeding CMV-STIL^+/-^ mice with animals expressing a dominant-negative version of p53 (p53-R172H; Supplemental Fig. S3) (Olive et al. 2004) did not rescue proliferation. Graded inhibition of proliferation and accumulation of cells in interphase explains why CMV-STIL^+/-^ and CMV-STIL^+/+^ MEFs contain increasing frequencies of micronuclei and aberrant karyotypes (Fig. 2) despite similar levels of supernumerary centrosomes (Fig. 1C,D).

**Figure 3.**
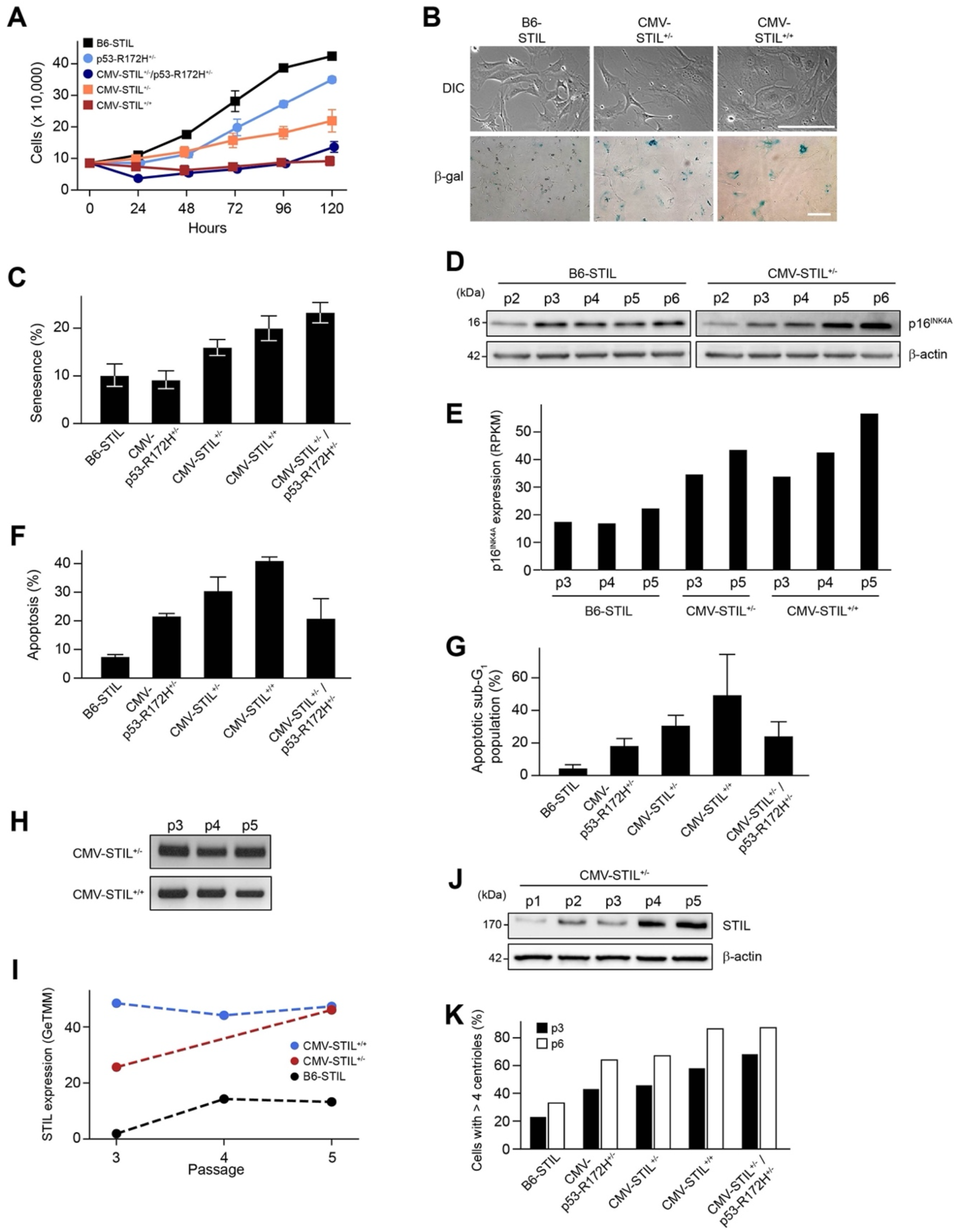
STIL overexpression impairs proliferation and induces senescence and apoptosis *in vitro*. (*A*) Proliferation of B6-STIL control, CMV-STIL^+/-^, CMV-STIL^+/+^, CMV-p53-R172H^+/-^ and CMV-STIL^+/-^/p53-R172H^+/-^ MEFs. Cells were enumerated every 24 h for a total of 5 days. Data are means ± SEM from three independent experiments. (*B*) Representative images of B6-STIL control, CMV-STIL^+/-^ and CMV-STIL^+/+^ MEFs, stained for β-galactosidase-cleaved x-Gal (lower panel). In the upper panel differential contrast interference (DIC) images of the cells are shown. Scale bar, 1 µm. (*C*) Percentage of β-galactosidase-positive, senescent B6-STIL control, CMV-STIL^+/-^, CMV-STIL^+/+^, CMV-p53-R172H^+/-^ and CMV-STIL^+/-^/p53-R172H^+/-^ MEFs. Data are means ± Clopper-Pearson 95%-CI. (*D*) Immunoblotting showing p16^INK4A^ protein expression levels in protein extracts from p2-6 CMV-STIL^+/-^ and B6-STIL control MEFs. β-actin served as a loading control. (*E*) RNA sequencing showing p16^INK4A^ mRNA levels in CMV-STIL^+/-^ and CMV-STIL^+/+^ relative to B6-STIL control MEFs. Percentage of (*F*) Apo-15-positive and (*G*) sub-G_1_ phase, apoptotic B6-STIL control, CMV-STIL^+/-^, CMV-STIL^+/+^, CMV-p53-R172H^+/-^ and CMV-STIL^+/-^/p53-R172H^+/-^ MEFs. Data are means ± SEM from three independent experiments. (*H*) Genotyping of CMV-STIL^+/-^ and CMV-STIL^+/+^ p3-5 MEFs. All passages harbor the STIL transgene. (*I*) RNA sequencing showing STIL mRNA levels in p3-5 B6-STIL control, CMV-STIL^+/-^ and CMV-STIL^+/+^ MEFs. (*J*) Immunoblotting showing STIL protein expression levels in p1-5 of CMV-STIL^+/-^ MEFs. β-actin served as a loading control. (*K*) Percentage of p3 and p6 B6-STIL control, CMV-STIL^+/-^, CMV-STIL^+/+^, CMV-p53-R172H^+/-^ and CMV-STIL^+/-^/p53-R172H^+/-^ MEFs with supernumerary centrioles.

CMV-STIL^+/-^ and CMV-STIL^+/+^ MEFs showed increasingly enlarged, flattened and irregular shapes, suggestive of cellular senescence (Fig. 3B). As both centrosome loss and amplification as well as chromosome missegregation can trigger a senescence-like state (Santaguida et al. 2017; Arnandis et al. 2018) we determined whether cellular senescence contributes to the reduced proliferation of STIL-overexpressing MEFs. Indeed, STIL overexpression led to a graded increase of β-galactosidase-positive, senescent CMV-STIL^+/-^ and CMV-STIL^+/+^ MEFs (Fig. 3B,C). CMV-STIL^+/-^ MEFs showed early accumulation of the senescence marker p16^INK4A^ at both RNA and protein levels (Fig. 3D,E) (Lopez-Otin et al. 2013). Analogous to impaired proliferation, senescence was not rescued by co-expression of dominant-negative p53-R172H. Due to their severely impaired proliferation, insufficient numbers of CMV-STIL^+/+^ MEFs were available for immunoblotting.

Next, we assessed whether overexpression of STIL induces apoptosis, as shown for PLK4 *in vitro* and *in vivo* (Holland et al. 2012; Marthiens et al. 2013; Sercin et al. 2016). Fluorescence-activated cell sorting (FACS) analysis after labelling with Apo-15 peptide and 7-amino-actinomycin (7-AAD) demonstrated a marked increase in apoptosis in CMV-STIL^+/-^ and CMV-STIL^+/+^ MEFs, which could only be partially rescued by expression of p53-R172H in a CMV-STIL^+/-^ background (Fig. 3F). In line, FACS-based cell cycle analysis after propidium iodide staining showed a graded increase of the sub-G_1_ population in CMV-STIL^+/-^ and CMV-STIL^+/+^ MEFs (Fig. 3G).

STIL transgene as well as STIL mRNA and protein levels were maintained with passaging in consecutive early passages of CMV-STIL^+/-^ and CMV-STIL^+/+^ MEFs (Fig. 3H-J). Again, insufficient numbers of CMV-STIL^+/+^ MEFs were available for STIL immunoblotting. In line with continued STIL expression, the percentage of CMV-STIL^+/-^ and CMV-STIL^+/+^ cells with supernumerary centrosomes further increased with passaging, reaching about 80% for p6 CMV-STIL^+/+^ MEFs (Fig. 3K). Expectedly, interference with p53 function did not prevent STIL-induced centrosome amplification, as inactivation of p53 leads to supernumerary centrosomes in MEFs itself (Caulin et al. 2007).

Together, our data show that constitutive overexpression of STIL inhibits proliferation via stimulation of p53-independent senescence as well as p53-dependent and p53-independent apoptosis in a dose-dependent manner.

### STIL overexpression skews Mendelian inheritance, causes microcephaly and perinatal lethality, and shortens lifespan in mice

The mating of B6-STIL transgenic animals with CMV-CRE mice revealed a significant deviation of genotype distribution from Mendelian inheritance in weaned pups. Relative frequencies of both live-born CMV-STIL^+/-^ and CMV-STIL^+/+^ mice were significantly reduced for the benefit of B6-STIL control animals (Fig. 4A). Simultaneously, we found an increased frequency of pups that died around birth. Genotyping of still-born animals revealed that 8% belonged to the CMV-STIL^+/-^ and 92% to the CMV-STIL^+/+^ group. Similarly, mice overexpressing PLK4 in all epiblast-derived tissues during embryonic development have been described to die shortly after birth (Vitre et al. 2015). Also, litter sizes were reduced in mice with inducible PLK4 overexpression (Coelho et al. 2015) and common genomic variants spanning the PLK4 gene are associated with pregnancy loss in humans (McCoy et al. 2015).

**Figure 4.**
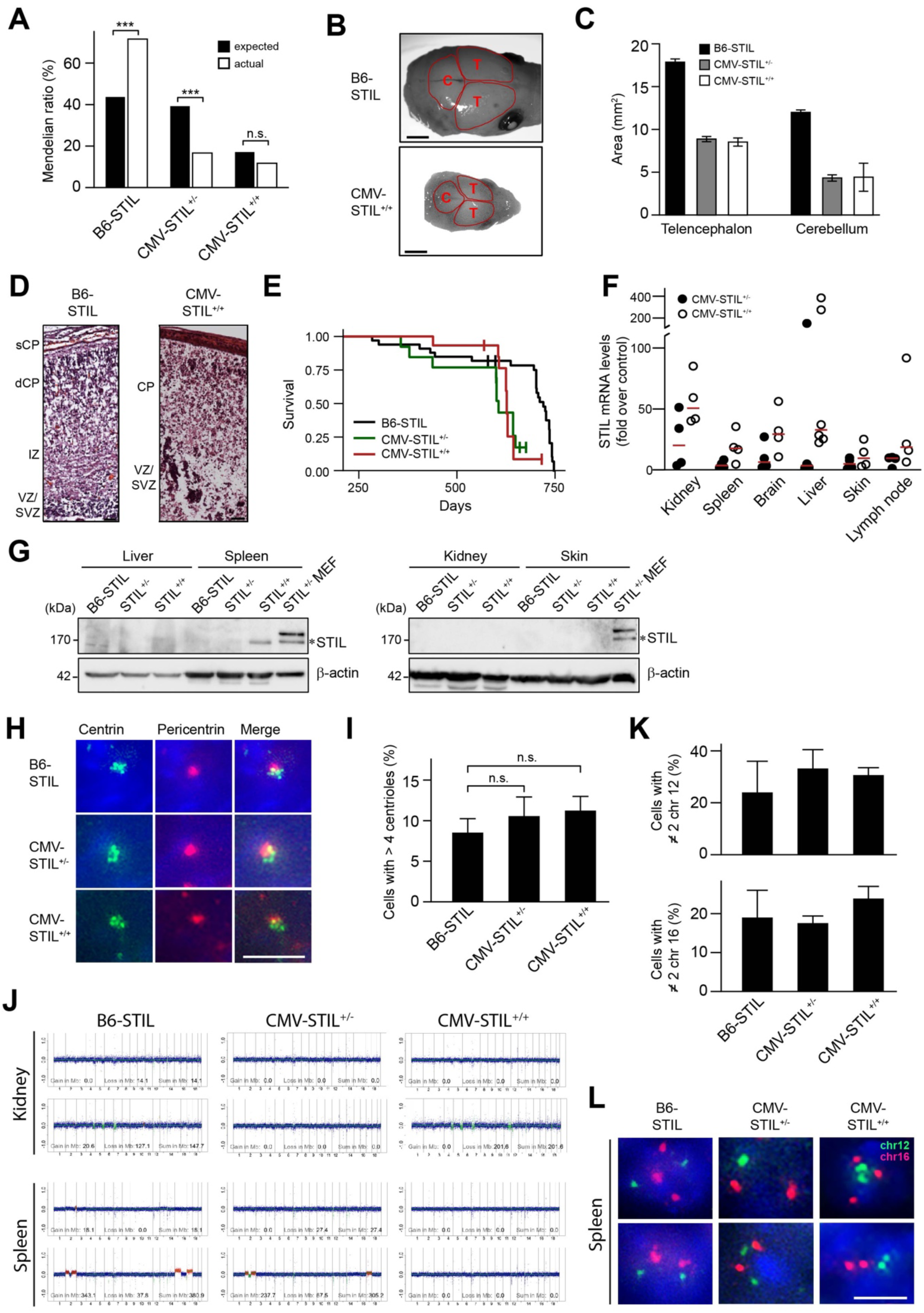
STIL overexpression skews Mendelian inheritance, causes microcephaly and perinatal lethality, and shortens lifespan in mice. (*A*) The graph shows the proportion of animals expected and obtained with the indicated genotypes. (*B*) Dorsal views of control B6-STIL and CMV-STIL^+/+^ brains at birth (P0). Telencephalic and cerebellar areas are encircled. Scale bar, 3 mm. (*C*) Mean telencephalic and cerebellar areas of the indicated genotypes at birth (P0) are depicted. Fifteen B6-STIL, three CMV-STIL^+/-^ and two CMV-STIL^+/+^ embryos were analyzed. Data are means ± SEM. (*D*) STIL overexpression disrupts cerebral cortical morphogenesis and organization. Representative H&E-stained sections of B6-STIL (left panel) and CMV-STIL^+/+^ (right panel) brains; Scale bars, 50 µm. (*E*) Overall survival of 30 B6-STIL, 10 CMV-STIL^+/-^ and 11 CMV-STIL^+/+^ animals. (*F*) Fold increase in STIL mRNA in tissues from CMV-STIL^+/-^ and CMV-STIL^+/+^ mice. Data are means ± SEM from three to six mice. Values were normalized to the respective B6-STIL tissue using the ΔΔCT method after normalization over HPRT. (*G*) Immunoblotting showing STIL protein expression levels in different organs from mice of the indicated genotypes. CMV-STIL^+/-^ MEF lysates were loaded as positive control. The STIL band at 170 kDa is marked with an asterisk. β-actin served as a loading control. (*H*) Representative images of centrosomes in splenocytes from B6-STIL (6 centrioles per centrosome), CMV-STIL^+/-^ (5 centrioles per centrosome) and CMV-STIL^+/+^ (6 centrioles per centrosome) mice. Cells were immunostained with centrin and pericentrin. Scale bar, 2 µm. (*I*) Percentage of cells containing >4 centrioles in spleens of B6-STIL, CMV-STIL^+/-^ and CMV-STIL^+/+^ mice. Data are means ± SEM from three mice per genotype. (*J*) WGS plots of healthy kidney and spleen tissues from mice with the indicated genotypes. Chromosomal region gains (red), losses (green) and the sums thereof in Mb are given for each plot. (*K*) Percentage of splenocytes with ≠ 2 dots of chromosome 12 (upper panel) and chromosome 16 (lower panel) per cell. Data are means ± SEM from three mice per genotype. (*L*) FISH using probes against chromosomes 12 (green) and 16 (red) performed on sections of healthy spleens from 2-year old B6-STIL, CMV-STIL^+/-^ and CMV-STIL^+/+^ mice. Nuclei were stained with DAPI. Scale bar, 5 µm. *p≤0.05, **p≤0.01, ***p≤0.001; n.s., not significant. sCP, superficial cortical plate; VZ/SVZ, ventricular/subventricular zone; IZ, intermediate zone; dCP, deep cortical plate.

Analogous to PLK4-overexpressing mice as well (Marthiens et al. 2013; Vitre et al. 2015), still-born CMV-STIL^+/-^ and CMV-STIL^+/+^ embryos showed a reduction of their brain size at birth (Fig. 4B,C). In line, germline truncating STIL mutations that lead to centrosome amplification cause primary microcephaly in humans (Kumar et al. 2009). To assess the effects of STIL overexpression on brain organization, we analyzed H&E-stained sections of B6-STIL control and CMV-STIL^+/+^ mice on postnatal day 0 (P0). Compared to B6-STIL control mice, the cerebral cortices of CMV-STIL^+/+^ mice were thinner, and lacked the typical laminar organization (Fig. 4D; Supplemental Fig. S4). Serial sectioning through the anterior/posterior extent of the brain failed to reveal clearly defined lateral ventricles in CMV-STIL^+/+^ animals (Supplemental Fig. S5).

Importantly, the median survival of CMV-STIL^+/-^ (606 days) and CMV-STIL^+/+^ mice (627 days) surviving to adulthood was significantly reduced compared to B6-STIL control animals (720 days) (*p*<0.005, logrank test), which translates into an approximately 15% decrease in lifespan (Fig. 4E).

To determine the levels of STIL overexpression and centrosome amplification *in vivo*, aged CMV-STIL^+/-^, CMV-STIL^+/+^ and B6-STIL control mice were sacrificed. In line with the results in MEFs, expression analysis by qPCR revealed a graded increase of STIL mRNA levels in all tissues analyzed in CMV-STIL^+/-^ and CMV-STIL^+/+^ animals (Fig. 4F). Lowest median STIL overexpression levels were found in skin and lymph nodes, and highest levels in kidneys, brain and liver, respectively. Analogous results were obtained by RNA sequencing (Supplemental Fig. S6). In contrast, immunoblotting revealed that STIL protein expression was not detectable in adult, aged mice, with the exception of spleen tissue from CMV-STIL^+/+^ animals (Fig. 4G), consistent with a translational shut down of the STIL transgene (Spriggs et al. 2010). Accordingly, even in spleens immunofluorescence analysis showed only a slight, not statistically significant increased frequency of cells containing supernumerary centrosomes in aged CMV-STIL^+/-^ and CMV-STIL^+/+^ mice (Fig. 4H,I).

To evaluate whether STIL overexpression causes clonal chromosomal aberrations *in vivo*, we performed whole genome sequencing (WGS) of spleens and kidneys from B6-STIL control and CMV-STIL mice with an age range from 20 to 104 weeks. No increased levels of chromosomal copy number alterations were found in CMV-STIL^+/-^ (mean gains + losses: 166.3 Mb; *p*=0.84) and CMV-STIL^+/+^ (mean gains + losses: 0.0 Mb; *p*=0.37) as compared to B6-STIL control spleens (mean gains + losses: 131.5 Mb) (Fig. 4J). Similarly and although STIL mRNA levels were highest in kidneys from STIL-transgenic mice, chromosomal copy number alterations levels were not increased in CMV-STIL^+/-^ (mean gains + losses: 0.0 Mb; *p*=0.35) and CMV-STIL^+/+^ (mean gains + losses: 142.6 Mb; *p*=0.88) as compared to B6-STIL control kidneys (mean gains + losses: 94.5 Mb). To assess at the single cell level whether numerical aneuploidy is more frequent in CMV-STIL-transgenic mice, splenocytes from aged B6-STIL control, CMV-STIL^+/-^ and CMV-STIL^+/+^ animals were analyzed by interphase fluorescence *in situ* hybridization (FISH) using probes to chromosomes 12 and 16. Confirming the WGS results, FISH did not reveal increased levels of whole chromosome aberrations in CMV-STIL^+/-^ and CMV-STIL^+/+^ as compared to B6-STIL control spleens (Fig. 4K,L).

These results show that STIL overexpression causes embryonic and perinatal lethality as well as microcephaly. Mice surviving to adulthood have a decreased lifespan. In those animals, selective pressure has eliminated cells with supernumerary centrosomes and aneuploidy by translational shut down of STIL overexpression in most tissues.

### STIL overexpression reduces spontaneous tumor formation

The by far most common type of malignancy and cause of death in C57BL/6 mice is lymphoma. 44.3% of these animals develop lymphomas, with about 90% of all neoplasms having occurred in C57BL/6 mice by the age of 24 months (Brayton et al. 2012). To determine whether STIL-induced centrosome amplification contributes to tumorigenesis, tumor formation in B6-STIL control, CMV-STIL^+/-^ and CMV-STIL^+/+^ mice was monitored up to an age of 24 months. Except for one sarcoma, the only tumors that emerged in any of the animals were lymphomas. In agreement with the literature, 50.0% of control mice developed lymphomas. Compared to B6-STIL control animals both CMV-STIL^+/-^ and CMV-STIL^+/+^ mice developed fewer tumors (Fig. 5A,B). Although slightly reduced, the mean age at tumor detection in CMV-STIL^+/-^ (628 days, *p*=0.28, logrank test) and CMV-STIL^+/+^ mice (636 days, p=0.46, logrank test) was not significantly different to B6-STIL control animals (674 days) (Fig. 5C). As the median lifespan of CMV-STIL mice is reduced below the mean age of tumor diagnosis (Fig. 4E), we conclude that STIL-transgenic mice likely developed fewer tumors because they died from other reasons before having reached the typical age of tumor onset.

**Figure 5.**
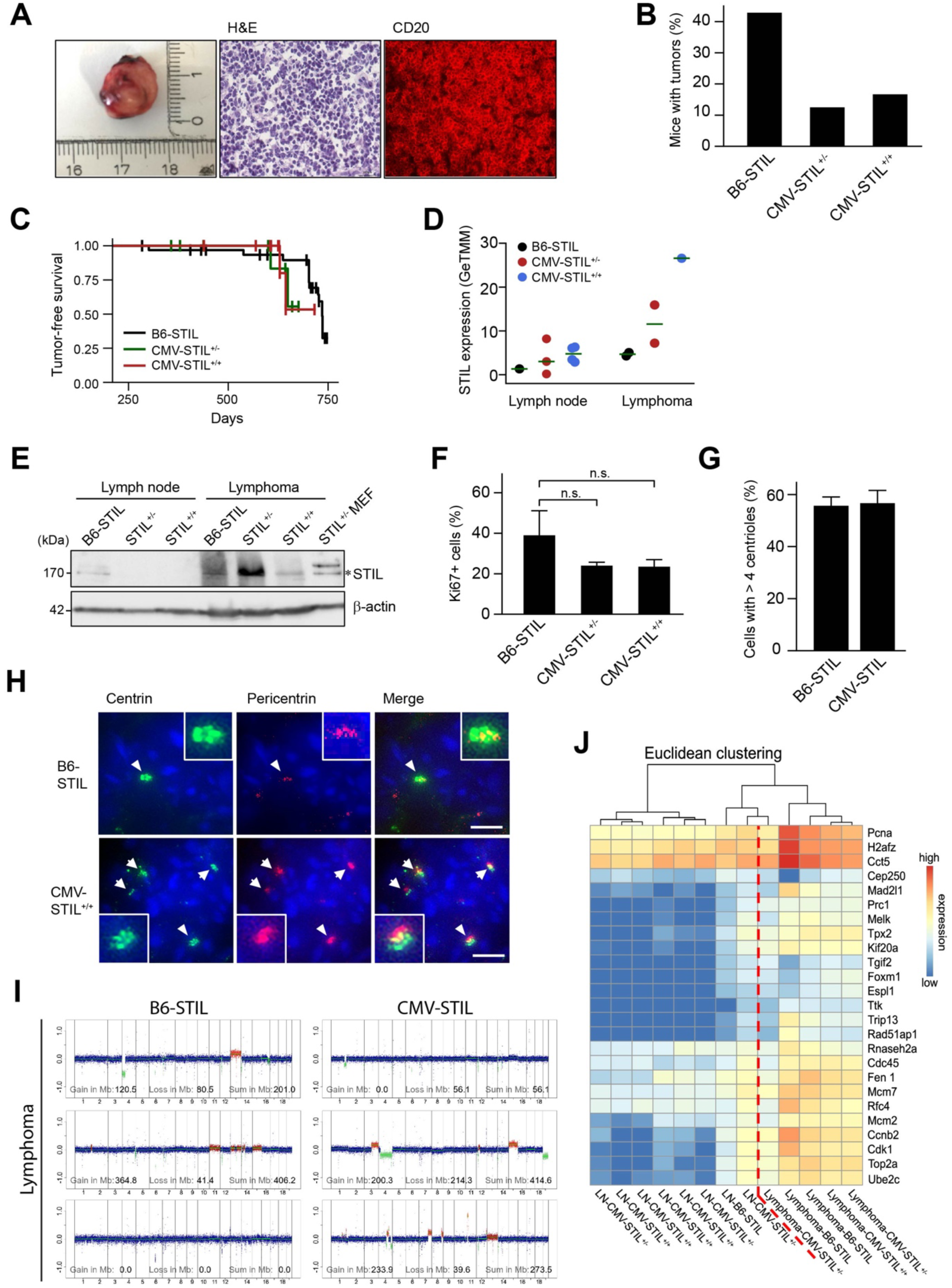
STIL overexpression shortens lifespan and reduces spontaneous tumor formation. (*A*) Macroscopy (left), histology stained with H&E (middle), and immunofluorescence stained with an antibody to CD20 (right) of a lymphoma arising from a B6-STIL mouse. (*B*) Percentage of mice with the indicated genotypes that developed tumors. (*C*) Tumor-free survival as quantified by the presence of macroscopic tumors at autopsy of 30 B6-STIL, 10 CMV-STIL^+/-^ and 11 CMV-STIL^+/+^ animals. (*D*) RNA sequencing showing STIL mRNA levels in normal lymph nodes and lymphomas from B6-STIL, CMV-STIL^+/-^ and CMV-STIL^+/+^ mice. (*E*) Immunoblotting showing STIL protein expression levels in normal lymph nodes and lymphomas from mice of the indicated genotypes. CMV-STIL^+/-^ MEF lysates were loaded as positive control. The STIL band at 170 kDa is marked with an asterisk. β-actin served as a loading control. (*F*) Percentage of Ki76-positive cells in healthy spleens and lymphomas from mice with the indicated genotypes. Data are means ± SEM from three mice per tissue and genotype. (*G*) Percentage of lymphoma cells from B6-STIL and CMV-STIL-transgenic mice with supernumerary centrioles. Data are means ± SEM from three B6-STIL versus two CMV-STL^+/-^ plus one CMV-STL^+/-^ lymphoma. (*H*) Immunofluorescence images of centrosomes in lymphomas from B6-STIL and CMV-STL^+/+^ mice, immunostained with antibodies to centrin and pericentrin. DNA is stained with DAPI (blue). Centrosomes are shown enlarged in insets. Scale bar, 5 µm. (*I*) WGS plots of lymphomas from mice with the indicated genotypes. Chromosomal region gains (red), losses (green) and the sums thereof in Mb are given for each plot. (*J*) Heat map showing normalized, RNA sequencing-derived expression levels of the CIN25 signature genes for healthy lymph nodes and lymphomas from B6-STIL, CMV-STL^+/-^ and CMV-STL^+/+^ mice. Samples were sorted by Euclidean clustering. The dashed red line separates normal lymph node from lymphoma samples. *p≤0.05, **p≤0.01, ***p≤0.001; n.s., not significant.

STIL mRNA levels gradually increased in lymphomas derived from B6-STIL control, CMV-STIL^+/-^ and CMV-STIL^+/+^ mice and were higher than the respective levels in healthy lymph nodes from B6-STIL control, CMV-STIL^+/-^ and CMV-STIL^+/+^ animals (Fig. 5D). Compared to healthy lymph nodes, STIL protein expression was increased in lymphomas from both B6-STIL control and CMV-STIL-transgenic mice irrespective of their genotype (Fig. 5E). STIL levels are low in early G_1_ phase and progressively increase until mitosis (Tang et al. 2011; Arquint et al. 2012; Vulprecht et al. 2012). Accordingly, STIL expression correlates with cellular proliferation and mitotic fraction of tissues, and is upregulated in multiple cancer types (Erez et al. 2004; Wang et al. 2022). In line, assessment of lymphomas from B6-STIL control, CMV-STIL^+/-^ and CMV-STIL^+/+^ mice by Ki67 immunostaining revealed that, corresponding to STIL protein levels, proliferation rates were elevated independent from lymphoma genotypes (Fig. 5F), suggesting that translational shutdown of STIL transgene expression has occurred in lymphomas as well and STIL protein expression is a consequence of increased lymphoma cell proliferation.

Immunofluorescence analysis showed that the frequency of cells harboring supernumerary centrioles was similar in B6-STIL control and CMV-STIL lymphomas, as 55.7 ± 5.9% (mean ± SEM) of B6-STIL control and 56.7 ± 8.5% of CMV-STIL-transgenic lymphoma cells exhibited centriole amplification (Fig. 5G,H). As overall only two CMV-STIL^+/-^ and two CMV-STIL^+/+^ mice developed lymphomas, results on supernumerary centrioles from those mice were pooled for this analysis.

To compare the degree of aneuploidy in lymphomas from B6-STIL control versus STIL-overexpressing mice, WGS of three lymphomas from CMV-STIL and B6-STIL control animals each was performed. In line with their similar levels of centrosome amplification, no statistically significant difference in the amounts of gained and/or lost base pairs between lymphomas derived from B6-STIL control (mean gains + losses: 202.4 Mb) versus CMV-STIL-transgenic mice (mean gains + losses: 249.1 Mb; *p*=0.79) was found (Fig. 5I). On the other hand, overall levels of chromosomal copy number aberrations were higher in lymphomas (mean gains + losses: 225.2 ± 173.7 Mb) as compared to healthy tissues (mean gains + losses: 87.3 ± 127.5 Mb; *p*=0.06), irrespective of their STIL transgene status (Fig. 4J; Fig. 5I), although the difference did not quite reach statistical significance. Accordingly, a chromosomal instability (CIN) score (Carter et al. 2006) inferred from RNA sequencing-derived gene expression profiles, separated healthy lymph node from lymphoma samples, irrespective of their STIL transgene status but was unable to discriminate between B6-STIL control, CMV-STIL^+/-^ and CMV-STIL^+/+^ samples (Fig. 5J).

Together, CMV-STIL^+/-^ and CMV-STIL^+/+^ mice were not predisposed to spontaneous tumor development but, instead, developed fewer lymphomas than B6-STIL control animals. This effect might be a consequence of the reduced lifespan of CMV-STIL^+/-^ and CMV-STIL^+/+^ mice due to reasons other than death from malignancies.

### STIL overexpression suppresses chemical skin carcinogenesis

Non-melanoma skin cancers represent the most frequent tumors in humans, with high levels of aneuploidy being associated with a poor prognosis in squamous skin carcinoma. Well established and standardized mouse models of squamous skin cancer recapitulate the features of human squamous skin carcinoma, with some degree of aneuploidy being present in all tumor cells (Abel et al. 2009), suggesting that squamous skin cancer represents an excellent model to study the role of supernumerary centrosomes and aneuploidy in tumor initiation.

To allow for centrosome amplification in mouse skin epidermis, we crossed B6-STIL control mice with animals expressing CRE under control of the keratin 14 (K14) promoter. As an analogous constitutive strategy led to a high rate of embryonic lethality in K14CRE-PLK4 mice (Sercin et al. 2016), we used CRE-ERT2, which encodes a fusion protein between CRE recombinase and the tamoxifen-responsive hormone-binding domain of the estrogen receptor, leading to CRE activation only after administration of tamoxifen (K14^CRE-ERT2^), similar to CRE-ERT2-driven overexpression of PLK4 described earlier (Vitre et al. 2015). The K14 promoter is active from early embryogenesis onwards in basal epidermal progenitor cells (Vasioukhin et al. 1999).

To determine whether STIL-induced centrosome amplification contributes to squamous skin carcinoma formation, K14^CRE-ERT2^-STIL^+/-^ mice were subjected to a classical two-stage skin carcinogenesis assay, in which a single topical application of a sub-carcinogenic dose of the chemical mutagen 7,12-dimethylbenz(a)anthracene (DMBA), which induces initiating *HRAS* mutations, is followed by multiple applications of the tumor promoter 12-O-tetradecanoyl-phorbol-13-acetate (TPA; three times weekly for 20 weeks) following a standardized protocol (Abel et al. 2009).

Intraperitoneal tamoxifen administration twice per week for two weeks to 35 ± 1 day old K14^CRE-ERT2^-STIL^+/-^ mice led to excision of the STOP cassette from the FLAG-STIL construct (Fig. 1A; Supplemental Fig. S7) and 5 to 10-fold increased expression of STIL mRNA in skin and esophagus tissue of K14^CRE-ERT2^-STIL^+/-^ animals, respectively (Fig. 6A,B). Analogous to K14^CRE^-PLK4 mouse embryos (Kulukian et al. 2015; Sercin et al. 2016), tamoxifen-induced STIL overexpression caused an about two-fold increase of centrin signals in both basal and suprabasal epidermal keratinocytes of K14^CRE-ERT2^-STIL^+/-^ animals (Fig. 6C,D).

**Figure 6.**
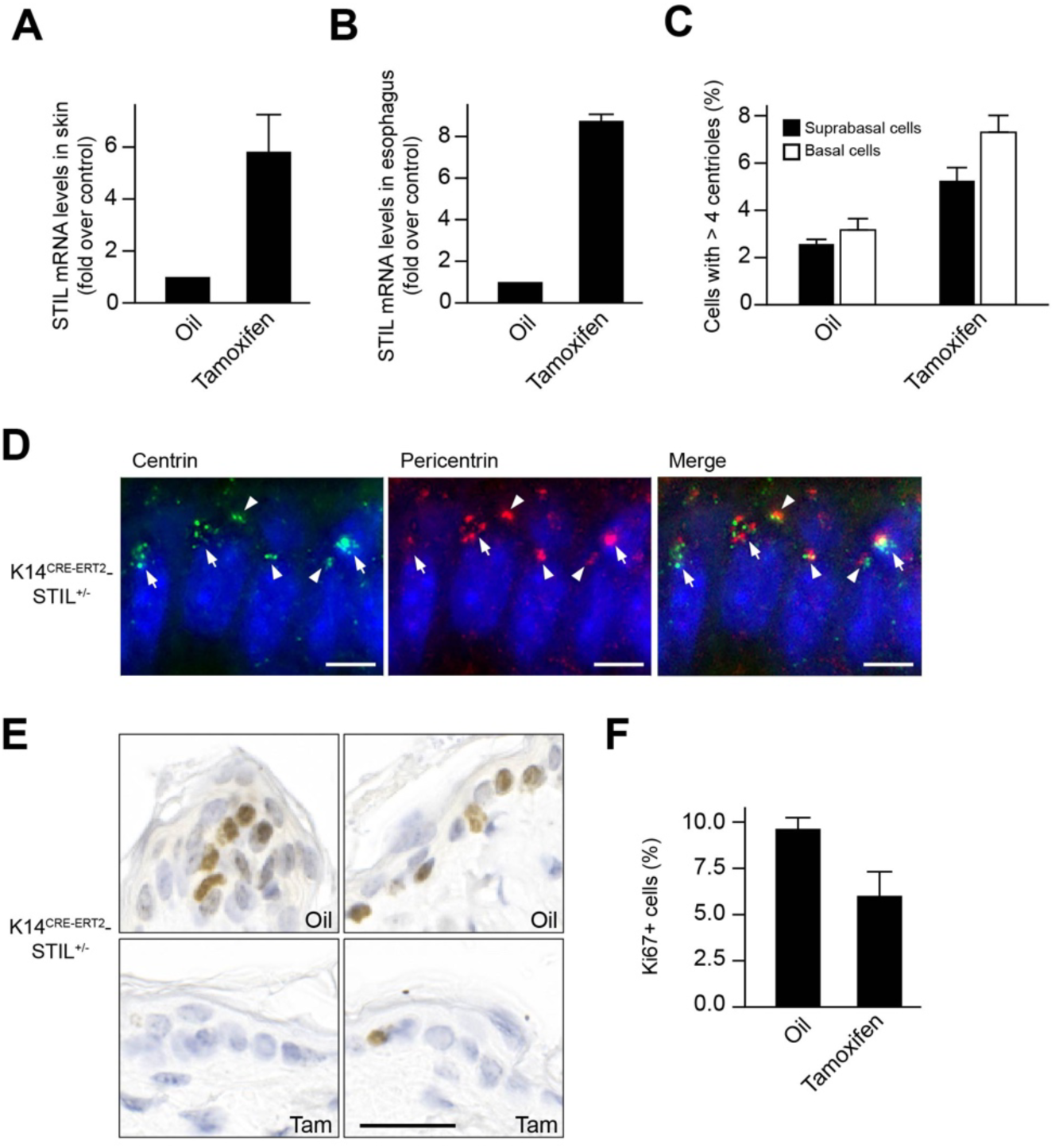
K14 promoter-driven STIL overexpression induces centrosome amplification and impairs proliferation in mouse skin. Quantitative RT-PCR showing the fold increase of STIL mRNA levels in (*A*) skin and (*B*) esophagus from tamoxifen-treated K14^CRE-ERT2^-STIL^+/-^ mice over oil-treated controls. Data are means ± SEM from four independent animals. (*C*) Representative immunofluorescence image of centrosomes in normal skin from a tamoxifen-treated K14^CRE-ERT2^-STIL^+/-^ mouse, immunostained with antibodies to centrin and pericentrin. DNA is stained with DAPI (blue). Centrosomes with regular and supernumerary centriole content are marked by arrow heads and arrows, respectively. Scale bar, 5 µm. (*D*) Percentages of basal and suprabasal epidermial cells from oil- versus tamoxifen-treated K14^CRE-ERT2^-STIL^+/-^ mice with supernumerary centrioles. Data are means ± SEM from three mice per condition. (*E*) Representative immunohistochemistry images of oil- versus tamoxifen-treated normal skin from K14^CRE-ERT2^-STIL^+/-^ mice, immunostained with an antibody to Ki67 (brown). Nuclei are counterstained with hematoxylin (blue). Scale bar, 25 µm. (*F*) Percentage of Ki67-positive epidermal cells from oil- versus tamoxifen-treated K14^CRE-ERT2^-STIL^+/-^ mice. Data are means ± SEM from three mice per condition. *p≤0.05, **p≤0.01, ***p≤0.001.

To determine the fraction of actively cycling cells, skin sections were immunostained with an antibody to Ki67. In line with our findings with CMV-STIL MEFs (Fig. 3A) and data from the skin of K14^CRE^-PLK4 mice (Kulukian et al. 2015; Sercin et al. 2016), the proportion of Ki67-positive cycling cells was lower in tamoxifen-treated K14^CRE-ERT2^-STIL^+/-^ mice than in oil-treated K14^CRE-ERT2^-STIL^+/-^ control animals one day after the last tamoxifen injection, although the difference did not quite reach statistical significance (*p*=0.06) (Fig. 6E,F).

The DMBA/TPA treatment protocol was applied to the back skin in cohorts of 63 tamoxifen-treated K14^CRE-ERT2^-STIL^+/-^ females and 32 oil-treated K14^CRE-ERT2^-STIL^+/-^ female control mice, and tumor development was followed over time. Tamoxifen-treated K14^CRE-ERT2^-STIL^+/-^ mice were less susceptible to skin carcinogenesis compared to oil-treated control animals (Fig. 7A,B). Whereas the median papilloma-free survival was 10.7 weeks in oil-treated K14^CRE-ERT2^-STIL^+/-^ control mice, tamoxifen-treated animals developed papillomas with a much longer latency (16.7 weeks; *p*<0.001, log-rank test) (Fig. 7A). In addition, the mean maximum number of papillomas per mouse was 4.0 in oil-treated controls but only 1.9 in tamoxifen-treated K14^CRE-ERT2^-STIL^+/-^ animals (*p*<0.001, Mann-Whitney-U test) (Fig. 7B).

**Figure 7.**
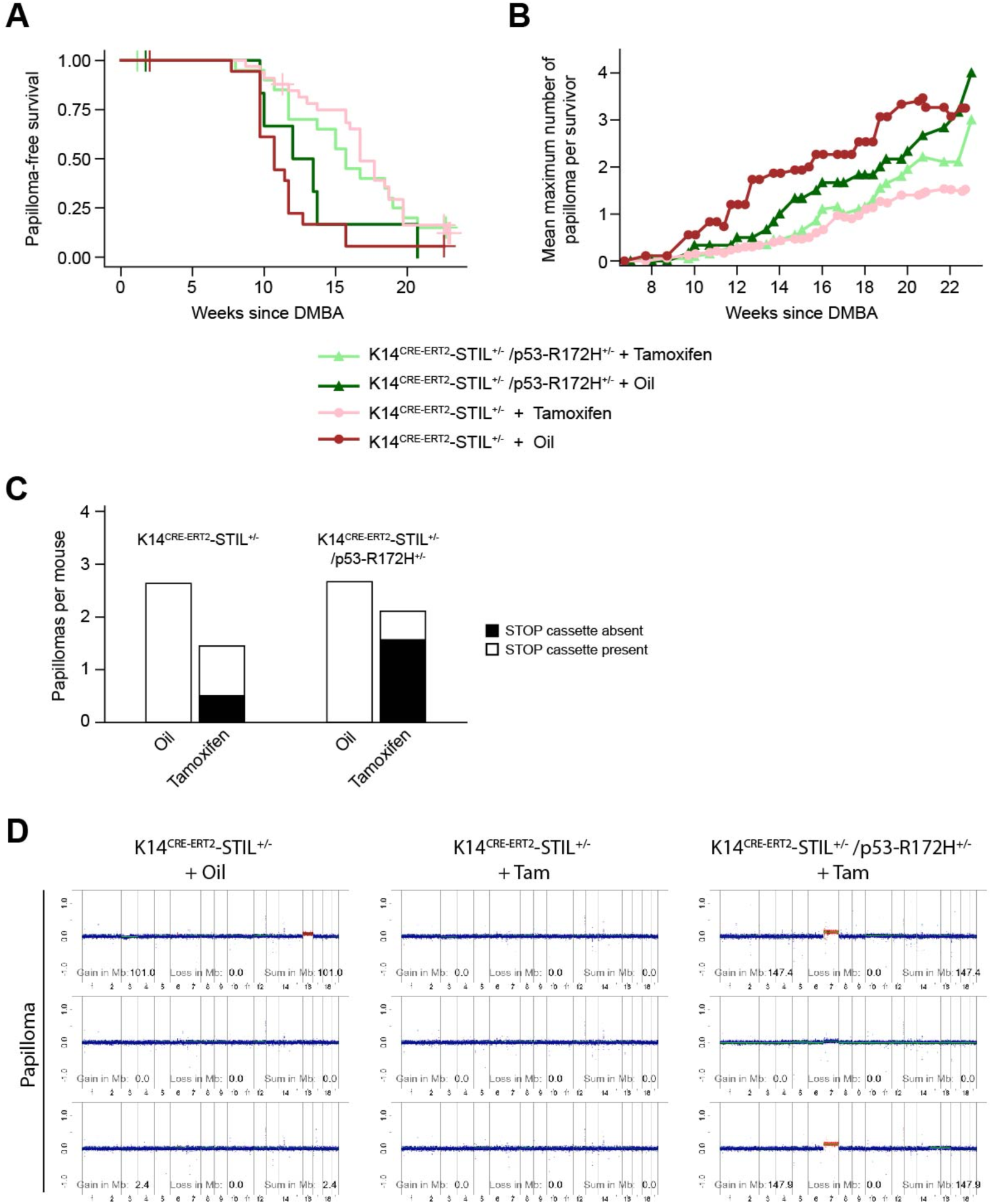
STIL overexpression suppresses chemical skin carcinogenesis. (*A*) Papilloma-free survival and (*B*) mean maximum number of papillomas per survivor as quantified by the presence of macroscopically visible skin tumors. Data result from the analysis of 14 oil- and 27 tamoxifen-treated K14^CRE-ERT2^-STIL^+/-^ mice, and 6 oil- versus 19 tamoxifen-treated K14^CRE-ERT2^-STIL^+/-^/p53-R172H^+/-^ animals over 25 weeks from the first tamoxifen treatment. (*C*) Genotyping for loxP-STOP-loxP cassette excision of 37, 39, 16, and 40 papillomas from oil- versus tamoxifen-treated K14^CRE-ERT2^-STIL^+/-^ and K14^CRE-ERT2^-STIL^+/-^/p53-R172H^+/-^ mice, respectively. (D) WGS plots of papillomas from mice with the indicated genotypes and treatments. Chromosomal region gains (red), losses (green) and the sums thereof in Mb are given for each plot.

Tamoxifen treatment causes CRE-ERT2-mediated excision of the STOP cassette and thereby induction of STIL expression in some but not all basal layer epidermal keratinocytes. Importantly, STOP cassette excision was found in only about one third of the papillomas from tamoxifen-treated K14^CRE-ERT2^-STIL^+/-^ mice, indicating that the majority of these papillomas originated from epithelial cells, which escaped activation of STIL transgene expression (Fig. 7C).

Several studies suggest that transient centrosome amplification induced by overexpression of PLK4 can accelerate tumorigenesis specifically in p53-deficient cells (Coelho et al. 2015; Sercin et al. 2016; Shoshani et al. 2021). To determine whether STIL- induced supernumerary centrosomes promote skin tumor formation in the absence of p53 function, we crossed K14^CRE-ERT2^-STIL^+/-^ animals with mice expressing dominant-negative p53-R172H. The p53-R172H missense mutation in mice corresponds to the p53-R175H hotspot mutation in human tumors and Li-Fraumeni syndrome. This mutation in the DNA binding domain results in a transcriptionally inactive protein that accumulates in cells, similar to the majority of naturally occurring versions of mutant p53 and therefore more faithfully recapitulates the situation in p53-mutant tumors than a p53 knockout (Yao et al. 2023). p53-R172H itself has been reported to increase skin tumor formation associated with centrosome amplification and induction of aneuploidy in a mouse skin carcinogenesis assay (Caulin et al. 2007). Nevertheless, papilloma-free survival was not significantly affected by concomitant p53 inactivation (*p*=0.68, log-rank test) (Fig. 7A). Similarly, mean maximum numbers of papillomas per mouse did not significantly differ between tamoxifen-treated K14^CRE-ERT2^-STIL^+/-^ and K14^CRE-ERT2^-STIL^+/-^/p53-R172H^+/-^ animals (*p*=0.22, Mann-Whitney-U test) (Fig. 7B). However, STOP cassette excision was increased to about two thirds of the papillomas in tamoxifen-treated K14^CRE-ERT2^-STIL^+/-^/p53-R172H^+/-^ animals (Fig. 7C).

The analysis of the genomic landscape of carcinogen-induced mouse skin tumors has shown that invasive squamous cell carcinomas, but not papillomas, present substantial chromosomal aberrations (Abel et al. 2009). Accordingly, WGS showed that papillomas from both oil- and tamoxifen-treated K14^CRE-ERT2^-STIL^+/-^ mice as well as K14^CRE-ERT2^-STIL^+/-^/p53-R172H^+/-^ animals were all largely diploid (Fig. 7D).

In conclusion, STIL overexpression suppresses DMBA/TPA-induced skin papilloma formation, an effect that is not rescued by concurrent expression of mutant p53.

## Discussion

Aneuploidy is a common characteristic of tumor cells and CIN is one of the hallmarks of cancer. However, in cultured cells including MEFs both aneuploidy and CIN often cause a proliferative disadvantage (Kops et al. 2004; Weaver et al. 2007; Funk et al. 2021). Accordingly, while low levels of aneuploidy and/or CIN, induced by interference with mitotic checkpoint proteins, are weakly tumor promoting in mice, high levels cause cell death and tumor suppression, and are associated with senescence and aging (Weaver et al. 2007; Lopez-Otin et al. 2013; Silk et al. 2013; Santaguida et al. 2017). Cell death induced by high CIN levels can occur through activation of p53, but p53 is not required for high CIN to suppress tumor formation (Kops et al. 2004; Thompson and Compton 2010; Santaguida et al. 2017; Funk et al. 2021). Complicating the interpretation of these findings, most proteins that function in mitosis and the mitotic checkpoint have additional, interphase roles outside of chromosome segregation, which often occur in pathways that are likely to influence tumor phenotypes.

One of the prime causes of aneuploidy and CIN is chromosome missegregation during mitosis induced by supernumerary centrosomes, which are a frequent finding in multiple tumor types (Gonczy 2015; Cosenza et al. 2017; Marteil et al. 2018). Although overexpression of PLK4 led to an elevated tumor incidence in mouse models (Coelho et al. 2015; Sercin et al. 2016; Levine et al. 2017; Shoshani et al. 2021), this causal relation has to be corroborated, as other models did not show elevated levels of spontaneous or carcinogen-induced tumor formation (Marthiens et al. 2013; Kulukian et al. 2015; Vitre et al. 2015). These discrepancies may, analogous to aneuploidy and/or CIN induction by interference with mitotic checkpoint proteins, be caused by differences in PLK4 expression levels or other roles of PLK4 in addition to centrosome replication.

To avoid effects outside the centrosome, we have generated a mouse model of centrosome amplification, in which the structural centriole protein STIL instead of PLK4 kinase is overexpressed (Tang et al. 2011; Arquint et al. 2012; Vulprecht et al. 2012). In these mice STIL overexpression is driven from a CMV promoter, which leads to transgene expression at levels similar to the CAG promoter used in most of the mouse models overexpressing PLK4 (Marthiens et al. 2013; Kulukian et al. 2015; Vitre et al. 2015; Sercin et al. 2016). We find that constitutive STIL overexpression causes centrosome amplification accompanied by aneuploidy, which however is highly deleterious for cell fitness even in the absence of p53 *in vitro*. Importantly, similar to reduced BubR1 levels and combined Bub3/Rae1 haploinsufficiency (Baker et al. 2004; Baker et al. 2006), graded overexpression of STIL caused increased senescence and apoptosis in MEFs. Supporting a dose effect of centrosome amplification similar to that observed for CIN, induction of excess multipolar spindles with subsequent chromosome missegregation seems to support paclitaxel cytotoxicity in breast cancer patients (Scribano et al. 2021). *In vivo*, in addition to induction of microcephaly, impaired skin stratification and perinatal lethality as previously reported for mice constitutively overexpressing PLK4 already (Marthiens et al. 2013; Coelho et al. 2015; Vitre et al. 2015; Sercin et al. 2016), constitutive overexpression of STIL led to a shortened lifespan. Again, these findings are reminiscent of BubR1-insufficient and Bub3/Rae1-haploinsufficient mice, for which signs of early aging and a shortened lifespan have been reported (Baker et al. 2004). Premature aging and reduced lifespan in mice with mitotic checkpoint gene defects are believed to be a consequence of cellular senescence (Baker et al. 2006). Interestingly, recent observations suggest that, in addition to DNA damage and oncogene activation, centrosome amplification can induce a senescence-like phenotype as well (Lopez-Otin et al. 2013; Arnandis et al. 2018; Wu et al. 2023).

CMV-STIL mice were not tumor prone. To the contrary, the frequency of tumors found in CMV-STIL animals was reduced compared to B6-STIL controls, possibly because STIL-transgenic animals had a shortened lifespan and therefore did not reach the median age of tumor onset. Similarly, BubR1-insufficient and Bub3/Rae1-haploinsufficient mice do neither show increased spontaneous tumor formation (Baker et al. 2004; Baker et al. 2006). Also, constitutive PLK4 overexpression from a CAG promoter in the brain only caused microcephaly instead of tissue overgrowth even in animals that additionally lacked p53 (Marthiens et al. 2013).

In mouse models with PLK4 overexpression, phenotypes seem to depend on both timing and levels of PLK4 expression. Transient, doxycycline-induced low level PLK4 overexpression in adult mice caused tumorigenesis both in the presence and absence of p53 (Coelho et al. 2015; Levine et al. 2017; Shoshani et al. 2021). On the other hand, constitutive overexpression of high PLK4 levels in mouse brain and skin from embryogenesis onwards causes tissue degeneration resulting in microcephaly (Marthiens et al. 2013) and impaired skin stratification (Kulukian et al. 2015; Sercin et al. 2016), respectively. Inactivation of p53 in addition to PLK4 overexpression led to further thinning of the skin and neuronal degeneration in two mouse models (Marthiens et al. 2013; Kulukian et al. 2015). In line with these findings, constitutive overexpression of STIL from embryonal development onwards caused microcephaly, perinatal lethality and reduced life span but did not increase spontaneous tumor formation either.

The mouse skin model of two-stage chemical carcinogenesis represents one of the best-established *in vivo* assays to evaluate the impact of genetic manipulation on tumor formation (Abel et al. 2009). To determine the role of transient STIL overexpression on skin tumorigenesis, we have therefore induced STIL expression in epidermal cells of K14^CRE-ERT2^-STIL^+/-^ mice by tamoxifen treatment and subsequently exposed their back skin to DMBA/TPA treatment. Despite STIL overexpression-induced centrosome amplification in both basal and suprabasal epidermal cells, papilloma formation was reduced in this system. Moreover, the majority of the papillomas in tamoxifen-induced, DMBA/TPA-treated K14^CRE-ERT2^-STIL^+/-^ mice originated from epithelial cells without CRE-mediated transgene recombination. Analogously, in mice with K14-CRE-driven constitutive overexpression of PLK4 in epidermal cells, DMBA/TPA treatment did not cause increased skin tumor formation and PLK4 -overexpressing mice even exhibited a trend toward lower overall skin tumor volume per animal (Vitre et al. 2015). We therefore suspect that TPA-driven proliferation of epidermal cells was impaired by expression of the STIL transgene, similar to the situation in CMV-STIL MEFs, and resulted in the gradual elimination of STIL-overexpressing epidermal cells with supernumerary centrosomes, thereby depleting the pool of cells available for transformation. In line, STIL transgene expression was translationally shut down in most adult tissues of STIL-transgenic mice. Also, in mice with K14-CRE-driven overexpression of PLK4 in the developing epidermis, cells with extra centrosomes were eliminated and replaced by cells in which the transgene has been transcriptionally turned off in the postnatal epidermis (Sercin et al. 2016). In this system, the transient presence of supernumerary centrosomes during embryogenesis led to the survival of few aneuploid cells, which eventually developed into skin tumors in a p53-deficient background.

Papilloma formation in DMBA/TPA-treated K14^CRE-ERT2^-STIL^+/-^ mice could not be rescued by concurrent expression of dominant-negative p53-R172H, a mutant p53 version that itself induces centrosome amplification, aneuploidy and skin tumor formation in a chemical skin carcinogenesis assay in mice (Caulin et al. 2007). Given the dose dependency of effects from both, CIN and centrosome amplification, it is not unexpected that p53-R172H did neither rescue proliferation of CMV-STIL MEFs nor papilloma formation.

In summary, both our *in vitro* and *in vivo* experiments demonstrate that powerful mechanisms lead to the elimination of cells with extra centrosomes and/or aneuploidy by impaired proliferation, senescence and apoptosis. Senescence- and apoptosis-driven depletion of the stem cell pool may explain reduced life span and tumor formation in STIL-transgenic mice. Shorter life span per se might contribute to the reduced tumor incidence in CMV-STIL mice as well, because these animals did not reach the median age of tumor onset in B6-STIL controls.

## Materials and methods

### Mouse lines

All animal studies were performed in accordance with the German Animal Protection Legislation, and were approved by the Institutional Animal Care and Use Committee and the Animal Care Committee of the Regierungspräsidium Karlsruhe, Germany (registration number: 35-9185.81/G-59/17). Mice were maintained under defined specified pathogen-free conditions according to FELASA regulations. Their health status was monitored daily. Three weeks-old mice were ear-marked and ear punches were genotyped as detailed below. If necessary for downstream applications, mice were sacrificed by cervical dislocation in compliance with the European guidelines for the care and use of laboratory animals.

B6-STIL (STIL^fl/fl^) mice were generated by cloning the Flag-tagged STIL cDNA construct into a Rosa26 targeting vector that contained neomycin resistance, loxP-STOP-loxP and IRES-mCherry sequences. This vector (Gt(ROSA)26Sor^tm1(Stil-IRES-mCherry)Pg^) was then transfected in C57BL/6 embryonic stem (ES) cells by electroporation. Neomycin-resistant cells were selected by PCR screening. Vector-positive C57BL/6 ES cell clones were subsequently injected into C57BL/6 blastocysts, which were transferred into a Black 6 (B6) foster mice. The resulting chimeric offspring was mated to verify germline transmission and bred for another three rounds.

B6-STIL mice were crossbred with CMV-Cre animals to generate CMV-STIL^+/-^ (CMV-Cre;STIL^lox/wt^) mice that overexpress murine STIL under the transcriptional control of the ubiquitous Rosa26 promoter after Cre-recombinase-mediated excision of an upstream STOP cassette. STIL^+/-^ mice were subsequently back-crossed to B6-STIL mice to obtain CMV-STIL^+/+^ animals. Mice were continuously examined and monitored for apparent developmental defects and spontaneous tumor development over a period of 24 months.

To generate mice with conditional STIL overexpression in K14-expressing epithelial cells, B6-STIL mice were crossbred with K14^CreERT2^ mice to obtain K14^CreERT2^-STIL^+/-^ animals. Conditional STIL overexpression with TP53 inactivation in K14-expressing epithelial cells was generated by breeding of K14^CreERT2^-STIL^+/-^ mice with P53-R172H^+/-^ mice to receive K14^CRE-ERT2^-STIL^+/-^/p53-R172H^+/-^ animals (Olive et al. 2004).

### Proliferation assay

Proliferation of MEFs (P3) was evaluated daily for 5 consecutive days by trypan blue staining with automated cell counting using a TC20™ counter (Bio-Rad). At day 0, 10^4^ cells were plated per 1 cm^2^ growth area of cell culture plate, each MEF line in triplicate.

### Senescence analysis

For senescence analysis, MEFs were plated in triplicate into 24-well plates. As positive and mock controls wildtype MEFs were treated with 100 nM paclitaxel and DMSO, respectively for 48 h. Thereafter, MEFs were stained using the Senescence β-Galactosidase Staining Kit (Cell Signaling) according to manufacturer’s instructions and eosin as cytoplasmic counter staining to detect senescent cells.

### Isolation of mouse embryonic fibroblasts

Mouse embryonic fibroblasts (MEFs) from B6-STIL, CMV-STIL^+/-^ and CMV-STIL^+/+^ mice were isolated from E12.5 mouse fetuses. Embryo heads were used for verifying of respective genotypes as detailed below, while MEFs were prepared from the bodies. In brief, bodies were digested in 0.25% trypsin/EDTA solution at 4°C overnight with gentle agitation. The resulting suspension was plated in DMEM/GlutaMAX (Gibco), supplemented with 10% FBS (Sigma), 1% penicillin/streptomycin (Gibco) and 1% MEM Non-Essential Amino Acids Solution (Gibco), and cultivated at 37°C in a 5% CO_2_ humidified incubator. Cells were passaged when reaching 70-80% confluence.

### Genotyping, quantitative polymerase chain reaction

For genotyping, DNA was isolated using the SampleIN Direct PCR Kit (highQu) or the AllPrep DNA/RNA/Protein Mini Kit (Qiagen), and amplified with ALLin HS Red Taq Mastermix (highQu) according to manufacturer’s instructions. Primer sequences used for genotyping are given in Supplemental Table S1. For quantitative polymerase chain reaction (qPCR), RNA from MEFs was isolated using the RNeasy Mini Kit (Qiagen). Mouse tissues were homogenized using TissueLyser II (Qiagen) and stainless-steel beads (5 mm, Qiagen) before RNA isolation using the AllPrep DNA/RNA/Protein Mini Kit (Qiagen) according to manufacturer’s instructions. RNA concentrations were measured using a NanoDrop™ 2000 Spectrophotometer (PeqLab). RNA was reverse transcribed using the QuantiTect Rev Transcription Kit (Qiagen) according to manufacturer’s instructions. qPCR was performed in 348-well plates on a LightCycler 480 instrument (Roche) using the Quantitect SYBR Green PCR Kit (Qiagen). All reactions were done in triplicate. Primer sequences used for qPCR are given in Supplemental Table S1. HPRT and PIPB were used as reference genes. Relative mRNA amounts were calculated using the comparative C_t_ method after normalization to reference gene expression.

### Whole genome sequencing and RNA sequencing

Genomic DNA and RNA from healthy tissues and lymphomas of control and STIL-transgenic mice was extracted using the All Prep DNA/RNA/Protein Mini Kit according to manufacturer’s instructions after tissue homogenization using TissueLyser II and stainless-steel beads (5 mm, all Qiagen). DNA and RNA concentrations were quantified with Qubit 2.0 using the dsDNA High Sensitivity and RNA High Sensitivity Kit (both Thermo Scientific), respectively. Whole genome sequencing (WGS) and RNA sequencing libraries were prepared using the Illumina TrueSeq Nano DNA Kit and TrueSeq Stranded RNA Kit, respectively, according to manufacturer’s instructions. Equimolar multiplexed libraries were then sequenced using 100 or 150 bp paired-end runs on HiSeq 4000 or NovaSeq 6000 S4 platforms (Illumina) at the DKFZ Genomics and Proteomics Core Facility. The resulting raw sequences were in FASTQ format, which were aligned to the mouse reference genome as bam files by the DKFZ OTP pipeline (Reisinger et al. 2017). We used the R-package HMMcopy (Ha et al. 2012) to call chromosome copy number variations from WGS data, as this method is applicable to inbred strains (Lange et al. 2020). To specify RNA expression levels, gene length corrected TMM (GeTMM)(Smid et al. 2018) values that allow for simultaneous intra- and inter-sample normalization were used.

### Immunoblotting

Cell lysis and immunoblotting were performed according to standard protocols. For immunoblotting, MEFs were lysed in RIPA buffer (50 mM Tris/HCl, pH 7.4, 1 mM NaCl, 1 mM EDTA, 0.25% sodium deoxycholate, 1% IGEPAL CA-630) supplemented with complete protease and PhosSTOP phosphatase inhibitor cocktails (Roche). Mouse tissues were mechanically homogenized by manual grinding in liquid nitrogen before lysis in RIPA buffer. Protein extracts were separated on 10% or 15% SDS-PAGE gels and transferred to nitrocellulose membrane (Bio-Rad). Membranes were incubated with the primary antibodies listed in Supplemental Table S2 overnight, and detected with an appropriate species-specific, horseradish peroxidase (HRP)-conjugated secondary antibody (1:10,000, Dianova) and chemiluminescence (Clarity Western ECL Substrate, Bio-Rad). The specificity of the STIL antibody was verified by siRNA and CMV-Flag-STIL overexpression experiments in MEFs (Supplemental Fig. S8).

### Histology and immunohistochemistry

Tissues cryopreserved in Tissue-Tek® O.C.T. Compound (Sakura Finetek) were cryo-sectioned using a CM1950 cryomicrotome (Leica Biosystems). 8-10 μm sections on SuperFrost Plus slides (Thermo Scientific) were fixed in 1% paraformaldehyde for 10 min, washed in H_2_O for 2 min, and stained with hematoxylin/eosin (H&E). Subsequently, slides were washed in H_2_O, followed by dehydration in 75%, 95% and 100% ethanol dilutions, respectively, incubated in xylene for 10 min, and mounted using Neo-Mount (Sigma). Alternatively, tissues were fixed for 24 h in paraformaldehyde, dehydrated in a series of 75%, 95% and 100% ethanol dilutions, respectively, incubated in xylene and embedded in paraffin. 5 μm sections, prepared using a HM 355 S microtome (Thermo Scientific), were H&E stained.

### Immunofluorescence

Immunofluorescence (IF) staining was performed as described previously (Cosenza et al. 2017). Briefly, MEFs grown on coverslips were washed in PBS, treated with 0.5% Triton X-100 in PHEM buffer (60 mM PIPES, 25 mM HEPES, 8 mM EGTA, 2 mMMgCl_2_, pH = 6.9), washed for 5 min in PHEM buffer and fixed in ice-cold methanol/acetone for 7 min. For IF of mouse tissues, 8-10 μm cryosections were stored at -80°C were thawed at room temperature for at least 1 h, washed in PBS, treated with 0.5% Triton X-100 in PHEM buffer, washed for 5 min in PHEM buffer and fixed in ice-cold methanol/acetone for 7 min or 4% paraformaldehyde for 15 min. Following fixation, slides or coverslips were rehydrated in PBS for 5 min, permeabilized in 0.1% Triton X-100/PBS, blocked in 10% normal goat serum in PBS for 20 min, and incubated with primary antibodies for 1 h. Primary antibodies used are listed in Supplemental Table S2. After primary antibody incubation, slides or coverslips were washed 3 times for 5 min in PBS and incubated with appropriate species-specific Alexa Fluor-conjugated secondary antibodies (Molecular Probes). DNA was counterstained with Hoechst 33342 (Invitrogen). Coverslips with MEFs were mounted in Vectashield antifade mounting medium (Vector Laboratories). For tissue sections, the Vector TrueVIEWT Autofluorescence Quenching Kit was applied to eliminate tissue autofluorescence.

### Multiplex fluorescence in situ hybridization

Multiplex fluorescence *in situ* hybridization (M-FISH) was performed as previously described (Geigl et al. 2006; Cazzola et al. 2019). Briefly, seven pools of flow-sorted mouse chromosome painting probes were amplified and combinatorial labeled using DEAC-, FITC-, Cy3, TexasRed, and Cy5-conjugated nucleotides and biotin-dUTP and digoxigenin-dUTP, respectively, by degenerative oligonucleotide primed (DOP)-PCR. Prior hybridization, metaphase spreads fixed on glass slides were digested with pepsin (0.5 mg/ml; Sigma) in 0.2N HCL (Roth) for 10 min at 37°C, washed in PBS, post-fixed in 1% formaldehyde, dehydrated with a degraded ethanol series and air dried. Slides were denatured in 70% formamide/1x SSC for 2 min at 72°C. Hybridization mixture containing combinatorial labeled painting probes, an excess of unlabeled cot1 DNA in 50% formamide, 2x SSC, and 15% dextran sulfate were denatured for 7 min at 75°C, preannealed for 20 min at 37°C, and hybridized to the denatured metaphase preparations. After 48 hours incubation at 37°C slides were washed at room temperature in 2x SSC for 3 times for 5 min, followed in 0.2x SSC/0.2% Tween-20 at 56°C for 2 times for 7 min. For indirect labeled probes, an immunofluorescence detection was carried out. Therefore, biotinylated probes were visualized using three layers of antibodies: streptavidin Alexa Fluor 750 conjugate (Invitrogen), biotinylated goat anti avidin (Vector) followed by a second streptavidin Alexa Fluor 750 conjugate (Invitrogen). Digoxigenin labeled probes were visualized using two layers of antibodies: rabbit anti digoxin (Sigma) followed by goat anti rabbit IgG Cy5.5 (Linaris). Slides were washed in between in 4x SSC/0.2% Tween-20, 3 times for 5 min, counterstained with 4.6-diamidino-2-phenylindole (DAPI) and covered with antifade solution.

### Interphase fluorescence in situ hybridization

Interphase fluorescence *in situ* hybridization (FISH) was performed as described (Cosenza et al. 2017). In brief, coverslips were washed in 2x SSC for 5 min and RNA was digested by RNase A incubation for 1 h at 37°C. After 3 washes in 2x SSC, coverslips were incubated with pepsin/HCl solution for 12 min, washed 2 times with PBS/MgCl_2_, fixed with 1% formaldehyde and dehydrated by sequential 3 min washes in 70%, 85% and 100% ethanol. Centromere-specific fluorescence-labeled probes were incubated on coverslips and DNA denaturation was performed for 5 min at 76°C. Subsequently, samples were left to hybridize overnight at 42°C. Then, excess probe was washed away by a 10 min wash in 2x SSC at 66°C, followed by 2 washes in 0.2x SSC for 7 min. Finally, coverslips were dipped in 0.4x SSC/0.2% Tween-20, counterstained with DAPI and mounted in Vectashield antifade mounting medium (Vector Laboratories).

### Microscopy

For H&E- and senescence stainings, images were acquired with a widefield Zeiss Axiophot microscope supplied with an AxioCam MRc5 color camera (Zeiss). For IF and FISH, image analysis was performed on a full motorized inverse Zeiss Cell Observer.Z1 microscope equipped with an ApoTome.2 module and an AxioCam MRm camera (Zeiss) ZEN 3.2 (Blue edition, Zeiss) and ImageJ (Fiji) software was used for image analysis. For M-FISH, images of metaphase spreads were captured for each fluorochrome using highly specific filter sets (Chroma technology, Brattleboro, VT) and a DM RXA epifluorescence microscope (Leica Microsystems, Bensheim, Germany) equipped with a Sensys CCD camera (Photometrics, Tucson, AZ). Camera and microscope were controlled by the Leica Q-FISH software and images were processed on the basis of the Leica MCK software and presented as multicolor karyograms (Leica Microsystems Imaging solutions, Cambridge, United Kingdom).

### Fluorescence-activated cell sorting

For cell cycle analysis, 2 x 10^5^ MEFs were fixed in ice-cold methanol, washed in PBS with 1% FBS (Sigma) followed by incubation in PBS with 30 μg/ml RNase A for 30 min at 37°C. Then, 1 μg/ml propidium iodide (Molecular Probes) was added for additional 10 min. Fluorescence-activated cell sorting (FACS) analysis was done using an Accuri C6 (BD Biosciences) device. 15,000 events were counted by FSC/SSC with doublet exclusion but without exclusion of debris and apoptotic cells. Cell cycle analysis was done using FlowJo software (BD Biosciences). For apoptosis analysis, MEFs were trypsinized and stained with Apotracker-Green (Apo-15 peptide) and 7-AAD according to manufacturer’s instructions (Biolegend) to detect apoptotic cells. As positive control wildtype MEFs were treated with 4% paraformaldehyde for 60 min on ice prior to analysis. FACS analysis was done using an Accuri C6 (BD Biosciences) device.

### Chemical skin carcinogenesis

Under license G59/17 of the local authorities (Regierungspräsidium Karlsruhe, Germany), a standardized skin carcinogenesis assay (Abel et al. 2009) was combined with preceding tamoxifen-induced, Cre^ERT2^-mediated recombination. In brief, freshly prepared tamoxifen (1 mg/100 µl 10% ethanol in sunflower seed oil) or vehicle (100 µl 10% ethanol in sunflower seed oil) only as a control was administered intraperitoneally to 5-week old mice, which were put on Altromin 1324 FF diet throughout the experiment. A total of four doses (every 3-4 days) were given within 2 weeks, which leads to maximal induction of Cre^ERT2^-driven recombination (Gallini et al. 2023). At the age of 7 weeks, a single dose of 7,12-dimethylbenz(a)anthracene (400 nMol DMBA/100 µl acetone) was applied epicutaneously to the shaved back skin of the mice. One week later, 12-O-tetradecanoylphorbol 13-acetate (TPA) treatment (three times per week for a total of 20 weeks) was started by epicutaneous administration of 10 nMol TPA/100 µl acetone to promote tumor development. Papillomas >1 mm in diameter were scored weekly. Two weeks after the end of TPA administration, mice were sacrificed for necropsy by cervical dislocation.

### Statistical analysis

Statistical analyses were performed using the appropriate applications from the Python package SciPy, except survival analyses, which were performed using lifelines. In cases where Clopper-Pearson intervals were used as a measure of uncertainty, the rates of the corresponding subtests are equal.

## Acknowledgements

This project was supported by a grant of the Deutsche Forschungsgemeinschaft (DFG) (A.K.: KR 1981/4-1). We thank Polygene Transgenetics (Rümlang, Switzerland) for generation of B6- STIL mice. We are grateful to the Central Animal Laboratory, German Cancer Research Center Heidelberg (DKFZ) for breeding and maintenance of mice, and the Genomics and Proteomics Core Facility, German Cancer Research Center Heidelberg (DKFZ) for providing library preparation, sequencing and preliminary data processing services.

*Author contributions*: A.K. and M.R.C. conceptualized the study. A.M., M.R.C., K. B., A.H., A.P., K.M.-D., A.J., and B.K. performed experiments. T.W. performed bioinformatic, statistical and computational analyses. T.H.-L. performed statistical analyses. A.K., M.R.C., K.M.-D., A.J. and B.K. reviewed and edited the manuscript with input from all co-authors. T.W. and B.K. prepared the figures. A.K. wrote the original draft manuscript and supervised the study.

