## Supplemental Figures and Tables for "STIL overexpression shortens lifespan and reduces tumor formation in mice"

### STIL-induced centrosome amplification suppresses tumorigenesis in mice

[1]Clinical Cooperation Unit Molecular Hematology/Oncology, German Cancer Research Center (DKFZ) and Department of Internal Medicine V, University of Heidelberg, Heidelberg, Germany. [2]Department of Pathology, Faculty of Veterinary Medicine, Zagazig University, Zagazig, Egypt. [3]Genome Biology Unit, European Molecular Biology Laboratory (EMBL), Heidelberg, Germany. [4]Schaller Research Group, [5]Core Facility Tumor Models, [6]Division of Biostatistics, German Cancer Research Center (DKFZ), Heidelberg, Germany. [7]Institute of Human Genetics, [8]Department of Internal Medicine V, University of Heidelberg, Heidelberg, Germany.

**Supplemental Figures**

**A**

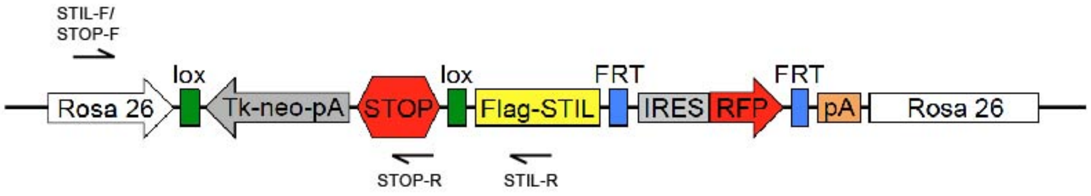


**B**

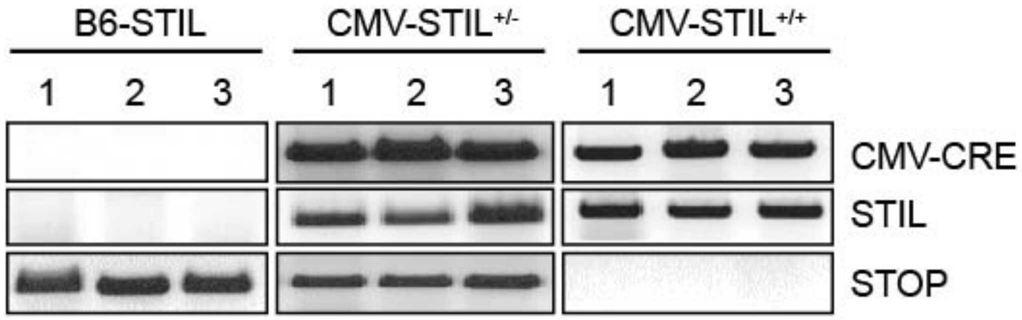


**Supplemental Figure S1.** STIL overexpression induces centrosome amplification and aberrant mitoses *in vitro*. (*A*) Schematic representation of the loxP-STOP-loxP-FLAG-STIL transgene. The localization of primers used for genotyping is indicated by arrows. (*B*) Genotyping of B6-STIL, CMV-STIL$^{+/-}$ and CMV-STIL$^{+/+}$ MEFs. For each MEF line three independent clones are shown. B6-STIL control MEFs lack CMV-CRE, whereas CMV-STIL$^{+/-}$ and CMV-STIL$^{+/+}$ MEFs are CMV-CRE positive. The FLAG-STIL transgene with excised loxP-STOP-loxP cassette is present in CMV-STIL$^{+/-}$ and CMV-STIL$^{+/+}$ but not B6-STIL MEFs. Biallelic loss of the loxP-STOP-loxP cassette in CMV-STIL$^{+/+}$ MEFs is verified by the absence of a STOP cassette PCR product.

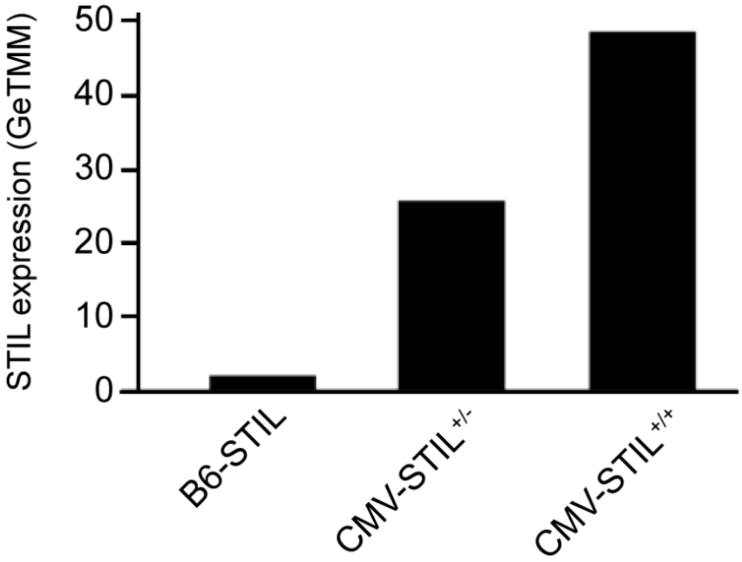


**Supplemental Figure S2.** STIL levels in MEFs by RNA sequencing. RNA sequencing showing STIL mRNA levels in MEFs (p3) from B6-STIL control, CMV-STL$^{+/-}$ and CMV-STL$^{+/+}$ mice.

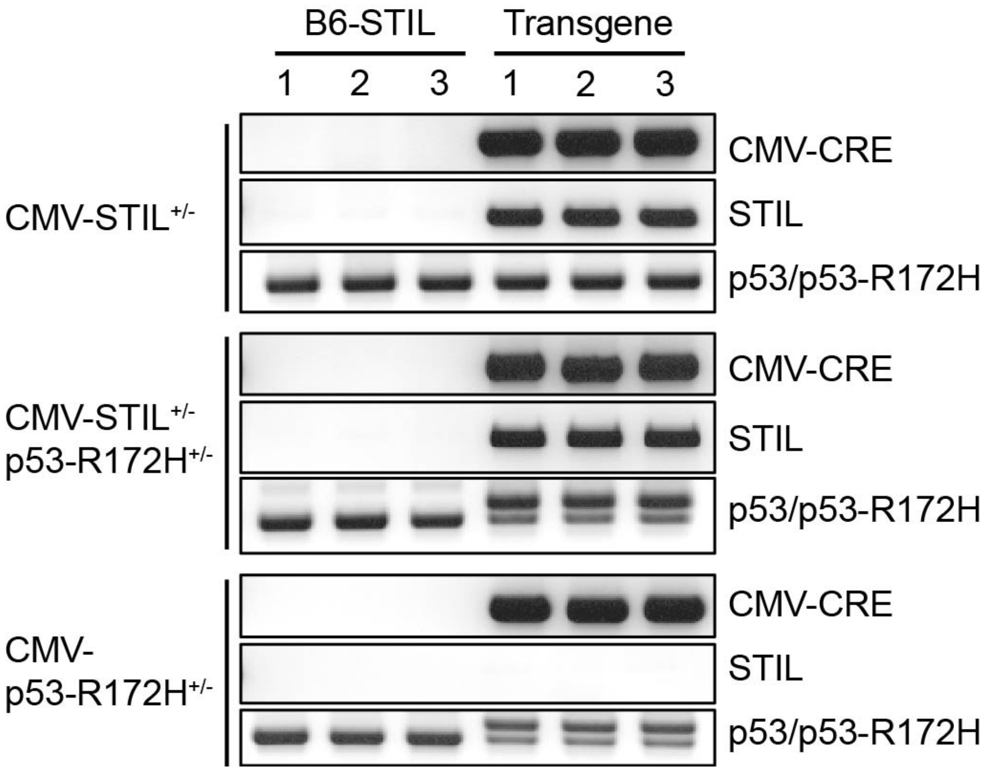


**Supplemental Figure S3.** STIL overexpression impairs proliferation and induces senescence and apoptosis *in vitro*. Genotyping of B6-STIL control, CMV-STIL$^{+/-}$, CMV-p53-R172H$^{+/-}$ and CMV-STIL$^{+/-}$/p53-R172H$^{+/-}$ MEFs. For each MEF line three independent clones are shown. B6-STIL control MEFs are negative for CMV-CRE and the STIL transgene with excised loxP-STOP-loxP cassette, and harbor only wildtype TP53. CMV-STIL$^{+/-}$ and CMV-p53-R172H$^{+/-}$ MEFs are both positive for CMV-CRE and the STIL transgene, and harbor only wildtype TP53 as well. CMV-p53-R172H$^{+/-}$ MEFs in addition harbor a mutant TP53-R172H allele (double band). CMV-p53-R172H$^{+/-}$ MEFs are positive for CMV-CRE but negative for the STIL transgene, and harbor both wildtype and mutant TP53.

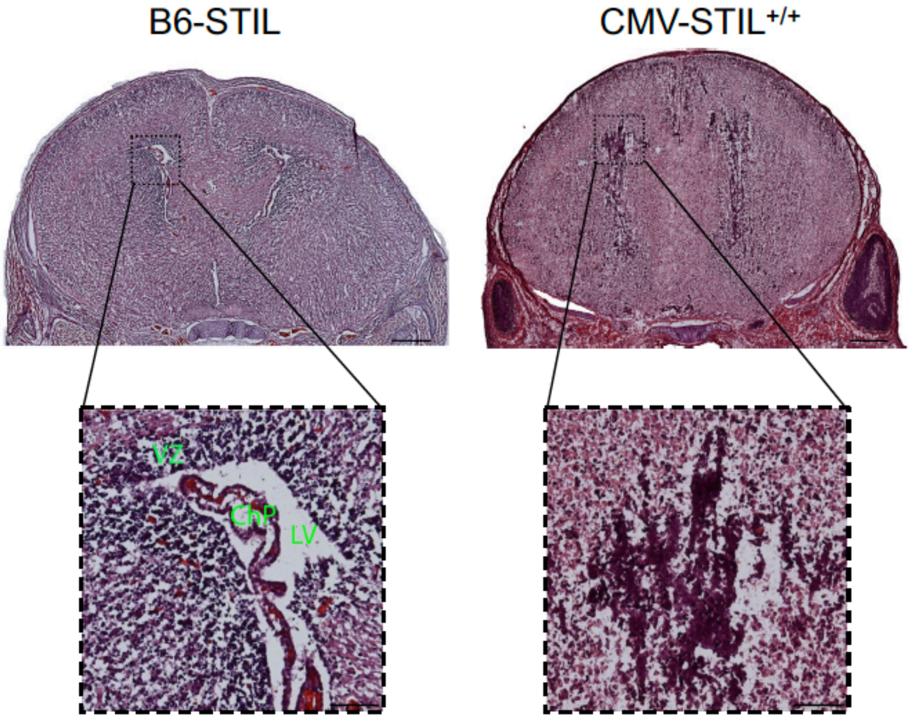


**Supplemental Figure S4.** Lack of lateral ventricles in CMV-STIL$^{+/+}$ mice. In contrast to brains from B6-STIL control mice at postnatal day 0 (left upper panel), the lateral ventricles (boxed regions in upper panels) appear to be collapsed in CMV-STIL$^{+/+}$ animals (right upper panel). The boxed regions in the upper panels are shown enlarged in insets (lower panels). VZ, ventricular zone; LV, lateral ventricle; ChP, choroid plexus. Scale bars in upper panels, 500 µm; Scale bars in insets, 100 µm.

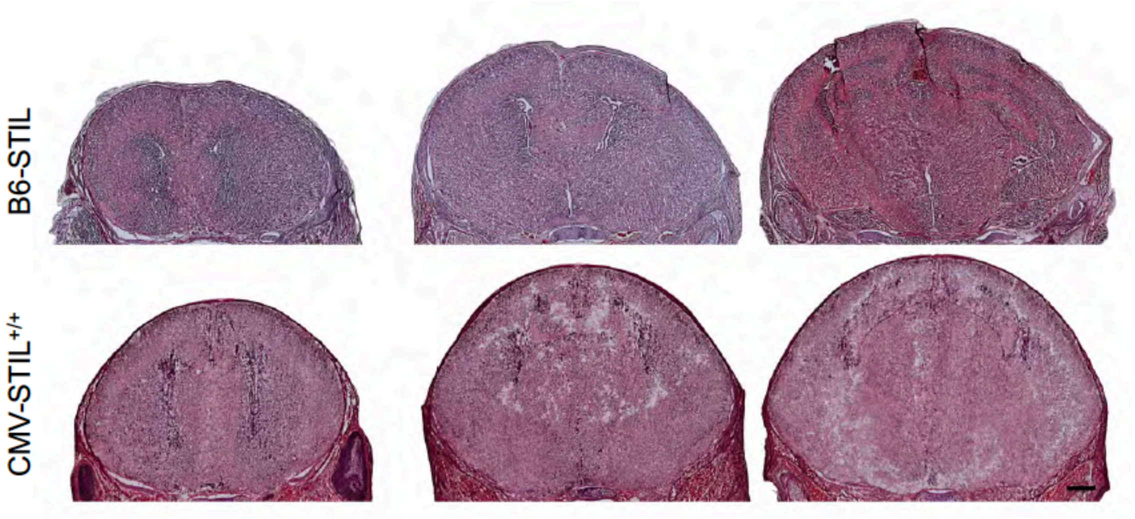


**Supplemental Figure S5.** Lack of lateral ventricles in CMV-STIL$^{+/+}$ mice. Serial sectioning through the anterior/posterior extent of the brain fails to reveal a clearly defined lateral ventricle in postnatal day 0 CMV-STIL$^{+/+}$ animals (lower panel). For comparison serial sections from a postnatal day 0 B6-STIL control mouse brain (upper panel) clearly depicting lateral ventricles are shown in the upper panel.

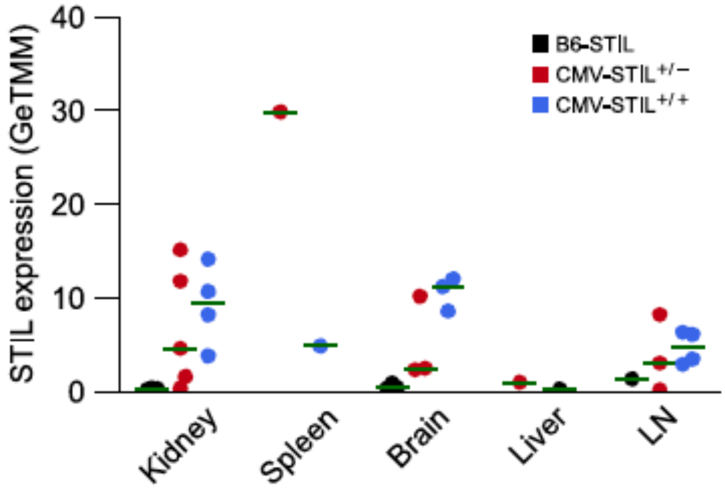


**Supplemental Figure S6.** STIL levels in healthy tissues by RNA sequencing. RNA sequencing showing STIL mRNA levels in different normal organs from B6-STIL control, CMV-STL$^{+/-}$ and CMV-STL$^{+/+}$ mice.

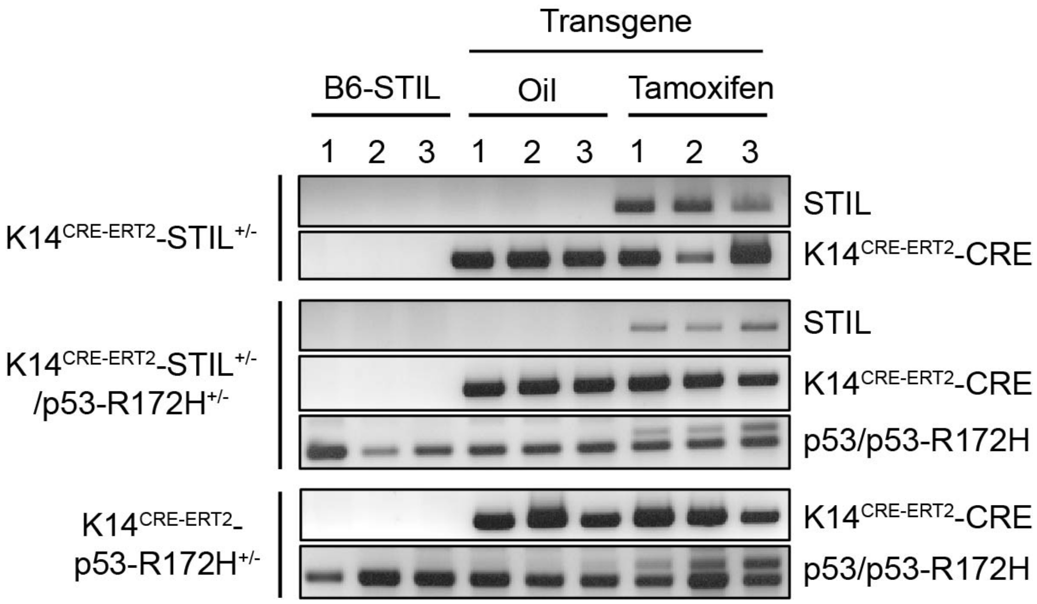


**Supplemental Figure S7.** K14 promoter-driven STIL overexpression induces centrosome amplification and impairs proliferation in mouse skin. Genotyping of B6-STIL, and oil- versus tamoxifen-treated K14$^{CRE-ERT2}$-STIL$^{+/-}$, K14$^{CRE-ERT2}$-STIL$^{+/-}$/p53-R172H$^{+/-}$ and K14$^{CRE-ERT2}$-p53-R172H$^{+/-}$ mice (n = 3 for each condition). The three B6-STIL mice are negative for K14$^{CRE-ERT2}$-CRE and the STIL transgene, and harbor only wildtype TP53. K14$^{CRE-ERT2}$-STIL$^{+/-}$ and K14$^{CRE-ERT2}$-STIL$^{+/-}$/p53-R172H$^{+/-}$ mice are positive for the STIL transgene, and K14$^{CRE-ERT2}$-STIL$^{+/-}$/p53-R172H$^{+/-}$ and K14$^{CRE-ERT2}$-p53-R172H$^{+/-}$ mice for mutant TP53 only after tamoxifen treatment.

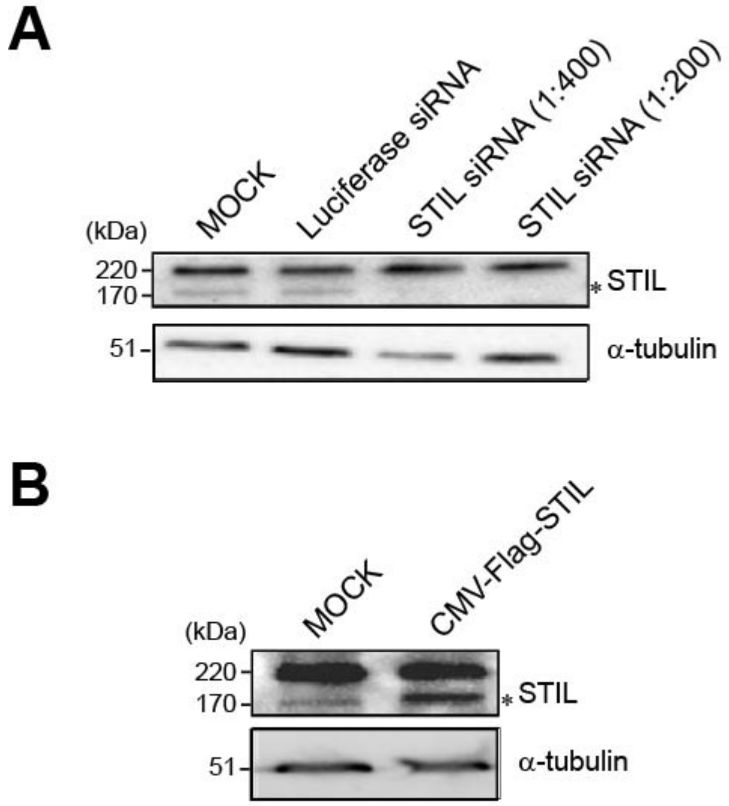


**Supplemental Figure S8.** Detection of mouse STIL protein in MEF lysates using the rabbit anti-STIL antibody A302-442A. (*A*) MEFs were transfected with control luciferase or STIL siRNA and immunoblotted with a rabbit anti-STIL antibody (A302-442A, Bethyl Laboratories) that detects human and mouse STIL at a size of 170 kDa (asterisk). siRNA-mediated knockdown of STIL led to the specific disappearance of the 170 kDa band. (*B*) Immunoblotting of lysates from MEFs transiently transfected with a CMV-Flag-STIL expression plasmid specifically enhanced the 170 kDa band when probed with the rabbit anti-STIL antibody (A302-442A, Bethyl Laboratories).

#### Supplemental Tables

**Supplemental Table S1.** Primers for genotyping and qPCR.

| | Name | Forward primer (5' - 3') | Reverse primer (5' - 3') |
|---|---|---|---|
| genotyping | STOP | AAAGTCGCTCTGAGTTGTTAT | GTGGCAGCTTCTTTAGCAAC |
| | STIL | AAAGTCGCTCTGAGTTGTTAT | CATCGTCGTCCTTGTAGTCAG |
| | CMV-CRE | GGCGCGGCAACACCATTTTT | CCGGGCTGCCACGACCAA |
| | K14-CRE | CGCCAATTAACCCTCACTAAAGG | ATCCATCAAATCGACCACCA |
| | TP53 | AGCCTGCCTAGCTTCCTCAGG | CTTGGAGACATAGCCACACTG |
| qPCR | STIL_qPCR | TCCTTGTGAGAGTAGGACGC | TCAAGGTCAGTGTCATGCTT |
| | HPRT | TGATCAGTCAACGGGGGACA | TTCGAGAGGTCCTTTTCACCA |
| | PBIB | TCGTCTTTGGACTCTTTGGAA | AGCGCTCACCATAGATGCTC |

**Supplemental Table S2.** Primary antibodies for immunoblotting. (IB) and immuno-fluorescence (IF).

| | Antibody | Clone | Company | Cat. No. | Dilution |
|---|---|---|---|---|---|
| IB | rabbit anti-STIL | | Bethyl Laboratories | A302-442A | 1:1000 |
| | mouse-β-actin-HRP | C4 | Santa Cruz | sc-47778 HRP | 1:5000 |
| | mouse-α-tubulin | DM1A | Sigma | T6199 | 1:5000 |
| | mouse-p16$^{INK4A}$ | F-4 | Santa Cruz | sc-74401 | 1:500 |
| IF | mouse-anti-centrin | 20H5 | Millipore | 1624 | 1:1000 |
| | rabbit-anti-pericentrin | | Abcam | ab4448 | 1:1000 |
| | rabbit-anti-Ki67 | D3B5 | Cell Signaling | 9129T | 1:100 |
| | rabbit-anti-CD20 | | Proteintech | | |
